# A single theory for the evolution of sex chromosomes and the two rules of speciation

**DOI:** 10.1101/2024.03.18.585601

**Authors:** Thomas Lenormand, Denis Roze

## Abstract

Sex chromosomes are involved in three major empirical patterns: Y (or W) chromosomes are often non-recombining and degenerate; heterogametic offspring (XY or ZW) from interspecific crosses are more often sterile or inviable than homogametic offspring (Haldane’s rule); the X (or Z) has a disproportionately large effect on reproductive isolation between species compared to autosomes (the large X effect). Each observation has received its own tailored explanation involving multiple genetic and evolutionary causes. Here, we show that these empirical patterns all emerge from a single theory for sex chromosome evolution incorporating the co-evolution of cis and trans-acting regulators of gene expression, and leading to systematic misexpression of dosage-compensated genes in heterogametic F1 hybrids, for both young and old sex chromosomes.

**Structured Abstract:** *Introduction:* Sex chromosomes have long been known to play a prominent role in the genetics of speciation, captured by the famous “two rules of speciation”. The first, attributed to J.B.S. Haldane and known as Haldane’s rule, corresponds to the general observation that when, among interspecific hybrids, one sex is inviable or sterile, it is more often the heterogametic sex (XY or ZW). The second is known as the large X effect, and refers to the fact that the X or Z chromosome often has a stronger effect on hybrid fitness than autosomes of equivalent size. While several theories have been proposed to explain these empirical patterns, none has been able to account for all the observations, leading to the current consensus that the two rules of speciation may have multiple causes. Furthermore, these theories remain disconnected from general models predicting sex chromosome evolution.

*Rationale:* We have recently developed a new theory for the evolution of sex chromosomes, showing that the coevolution of cis and trans regulators of gene expression can lead to stable recombination arrest between the X and Y (or Z and W) chromosomes, degeneration of the Y (or W) chromosome and dosage compensation. Here we investigate the role of sex chromosomes on the fitness of interspecific hybrids under this general model, considering different scenarios that correspond to different stages in the evolution of sex chromosomes. We also investigate the effect of genes with sex-specific effects that affect either male or female fertility.

*Results:* Haldane’s rule and the large X effect are observed in all scenarios considered. Our model also captures “Darwin’s corollary” to Haldane’s rule, i.e. an asymmetry between the effects of reciprocal crosses between species. These results are caused by the divergence of regulators of gene expression between species, due to the presence of a non-recombining, degenerate stratum on the Y chromosome either shared by both species or present in only one of them. In both cases, Haldane’s rule results from the heterogametic sex inheriting X chromosome cis regulators from only one parental species that have not coevolved with the trans regulators of the other species, resulting in under or overexpression of dosage-compensated genes on the X (or Z) chromosome. While this regulatory divergence may be limited in the case of species with a global (chromosome-wide) somatic system of dosage compensation, regulatory divergence of germline-expressed genes may generate Haldane’s rule for fertility in these species. Genes whose effect is restricted to the heterogametic sex often have strong effects on reproductive isolation and on the fitness of heterogametic hybrids, especially if they are silenced on the X (or Z) chromosome of one parental species and on the Y (or W) chromosome of the other.

*Conclusion:* Our study shows that a simple model of coevolution between cis and trans regulators of gene expression on sex chromosomes can explain the evolution of non-recombining, degenerate and dosage compensated Y and W chromosomes, Haldane’s rule and the large X effect. It may also explain why Haldane’s rule is based on viability is some taxa and on fertility in others, an observation that remained difficult to explain from previous theories. These results match with recent observations on the role of misregulation of gene expression on hybrid fitness, and open new perspectives of empirical research on the genetics of speciation.

Summary figure.
A multilocus, individual-based simulation model was used to investigate the joint evolution of sex chromosomes and reproductive isolation between species. It includes deleterious mutations, mutations suppressing or restoring recombination and mutations affecting the strength of cis and trans regulators of gene expression. This model predicts the major features of sex chromosome evolution and the “two rules of speciation” (Haldane’s rule and the large X effect).

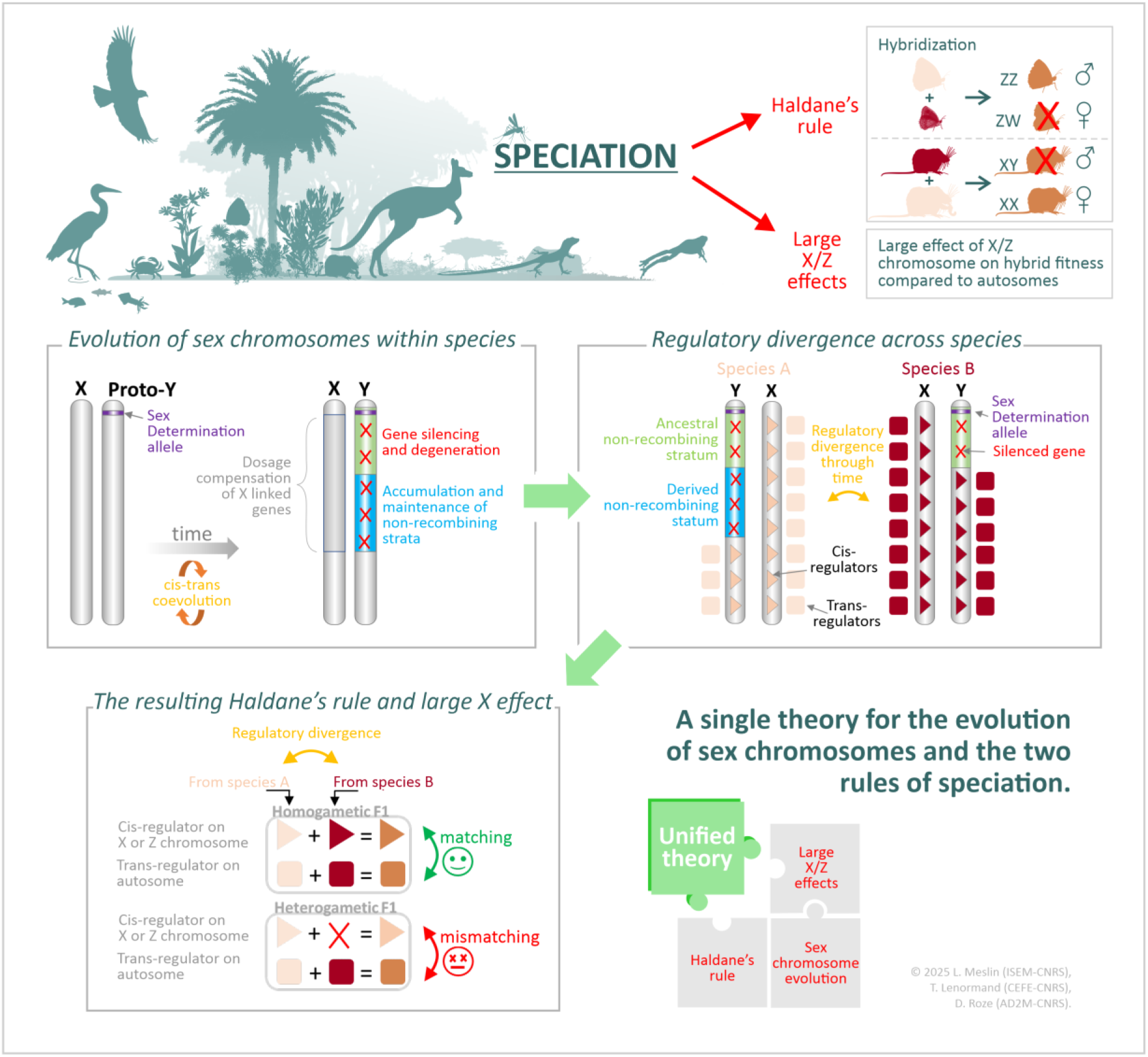

## Introduction

Sex chromosomes play a prominent role in the process of speciation, captured in the “two rules of speciation” (*1*). The first, ‘Haldane’s rule’ (HR), is named after the British scientist who observed that hybrid inviability or sterility occurs more frequently in the heterogametic than in the homogametic sex (*2*). Over a century of work has confirmed the generality of this observation in nature (*3–7*). Additionally, there is often an asymmetry of the effect observed between reciprocal crosses (*8*), termed ‘Darwin’s corollary’ to HR. The second, the ‘Large X effect’ (LX), refers to the observation that X chromosomes (and Z chromosomes in ZW species) disproportionately affect hybrid fitness compared with autosomes of equivalent size (*1*, *9–12*). These rules, among the few law-like generalizations in biology, have been extensively tested and studied, and have led to intense theoretical investigation to understand their origin (*3*, *5–7*, *13*, *14*).

Presently, the consensus is that HR and LX are composite phenomena with multiple genetic and evolutionary causes (*3*, *5–7*, *13–19*). While some theories have received more support than others, none offers a general solution—each failing to account for some observations. For example, a prominent explanation for the two rules is the dominance theory, first suggested by Muller (*20*) and later formalized (*21*, *18*, *22*), based on the idea that genetic incompatibilities involving at least one gene located on the X chromosome (or Z, in species with female heterogamety) may affect more strongly the fitness of the heterogametic sex if these incompatibilities are on average partially recessive. However, the theory does not explain why incompatibilities should be, on average, recessive (*21*, *23*), although fitness landscape models propose possible solutions (*24*–*26*, §3.2 in *27*). Furthermore, it does not explain well why HR often involves sterility rather than viability (as many more genes impact viability than fertility) (*16*, *28*), why it is observed in groups lacking a hemizygous X (*17*) or in groups where XX females only express one X, as in marsupials (*29*) or placentals (although in the latter case, both Xs may be expressed at the level of tissues (*18*)). Other theories better account for the importance of hybrid sterility in HR, but have their own major limitations. The “faster male theory” explains well why male sterility often occurs in hybrid crosses in some groups (*16*, *18*, *28*) and why it may occur in species lacking hemizygosity (*17*), but critically fails to account for HR in species where females are the heterogametic sex (*13*, *14*, *18*) (in these cases, a “faster female” theory would be required). The “meiotic drive theory” explains well why sex chromosomes could play a major role in the sterility of heterogametic hybrids (*30–32*). It has received some empirical support (*32–34*) but does not offer a convincing explanation for HR for viability, since the driving mechanism will only manifest itself when F1 hybrids undergo meiosis and attempt to produce gametes, but not before (*9*, *13*). Lastly, while sex chromosome degeneration is another general and intensely studied phenomenon, none of these theories considers that the processes leading to degenerate sex chromosomes and their subsequent evolution may be related to the emergence of either HR or LX (Table 1).

**Table. 1.** Summary of the comparison of main predictions of the different theories for Haldane’s rule and the large X effect. Regulatory theory refers to the theory presented in this paper. In green/orange/red the theory adequately/partially/inadequately predicts the pattern indicated in the first column. The cases where the theories fail or succeed are explained in the text. There are four “unsettled” cases (in orange and numbered): (1) To achieve full sterility, driver and anti-driver must be mutually incompatible, which should be revealed in both directions of the hybrid cross (*30*) (2) HR in mammal species with X inactivation in females could be explained by the dominance theory if HR concerns male sterility, but not male inviability. (3) Y-autosomes interactions are not considered in the usual dominance theory, as the argument is usually based on male hemizygosity resulting from Y degeneration by selective interference. However, as we show, male hemizygosity from X degeneration could evolve in a regulatory model for some male-limited genes. (4) The faster male theory, even if it does not specifically predict rare Y-autosome interactions, could accommodate this observation. (5) The observation of HR in species lacking a hemizygous X challenge all theories based on sex-chromosomes, but these species might actually have a small non-recombining region, sufficient to generate HR (see text). (6) The dominance theory postulates the existence of recessive incompatibilities, and the faster male theory postulates that males evolve incompatibilities more rapidly, but without providing a strong mechanistic or evolutionary explanation for why it should be so. Other less supported theories have been proposed, but not indicated on the table, see §3 in (*27*).

|  | *Domi-*<br>*nance*<br>*theory* | *Faster*<br>*male*<br>*theory* | *Meiotic*<br>*drive*<br>*theory* | *Regula-*<br>*tory*<br>*theory* |
| --- | --- | --- | --- | --- |
| HR in XY species | ✓ | ✓ | ✓ | ✓ |
| HR in ZW species | ✓ | ✗ | ✓ | ✓ |
| Large X/Z effect | ✓ | ✗ | ✓ | ✓ |
| Importance of HR through sterility in several groups | ✗ | ✓ | ✓ | ✓ |
| Importance of HR through inviability in several groups | ✓ | ✗ | ✗ | ✓ |
| Darwin's corollary to HR | ✓ | ✗ | (1) | ✓ |
| HR through inviability/sterility in species without/with global DC | ✗ | ✗ | ✗ | ✓ |
| HR in species with X inactivation in females | (2) | ✓ | ✓ | ✓ |
| HR on sterility sometimes caused by Y or W-autosomes interactions | (3) | (4) | ✓ | ✓ |
| HR on sterility in XY species with homomorphic sex chromosomes | (5) | ✓ | (5) | (5) |
| Mechanistic or evolutionary underlying explanation | (6) | (6) | ✓ | ✓ |
| Theory also explains evolution of sex chromosomes | ✗ | ✗ | ✗ | ✓ |

### Regulatory theory for sex chromosome evolution

We recently proposed a new theory for the evolution of non-recombining, degenerate, and dosage-compensated sex chromosomes based on XY regulatory divergence and the early emergence of dosage compensation (DC) (*35–37*). We show here that the same theory also predicts Haldane’s rule and the large X effect. In the present study, we followed the simulated independent evolution of 20 species using this previously described model (*35*) (Methods, §1.1 in (*27*), Fig 1). As detailed in the Methods and in (27), we performed individual-based stochastic simulations of populations of diploid individuals, with XY males and XX females (all the arguments below also apply to ZZ / ZW systems and large Z effect, but for simplicity we only discuss the XY case), incorporating deleterious mutations occurring at many genes, the evolution of recombination and the evolution of cis and trans regulation of gene expression (*35*). In this model, “lucky” Y inversions fix because they capture a portion of the Y with a lower-than-average load of deleterious mutations in the population (while also including the sex-determining locus), leading to recombination arrest between the X and Y chromosomes. Then, cis-regulatory divergence occurs together with an accumulation of deleterious mutations on the Y, and emerging gene-specific DC (*36*). Our model does not consider the evolution of global (i.e., chromosome wide) mechanisms of DC, but assumes that expression evolves on a gene-by-gene basis. The effect of global DC systems will be discussed later. This emerging DC creates sex-antagonistic effects that prevent the reestablishment of recombination, leading to a new degenerate, silenced and dosage compensated Y stratum. Once such a new stratum has evolved, cis- and trans-regulators continue to coevolve to maintain optimal expression in both males and females. We estimated the rate of occurrence and the pattern of hybrid incompatibilities by measuring the fitness of F1 hybrids among species at different time steps, under several scenarios of sex chromosomes at different stages of their evolution. We also compared these results to regulatory incompatibilities occurring on an autosome of equal size, in order to investigate the large X effect (see Methods).

**Fig. 1.**
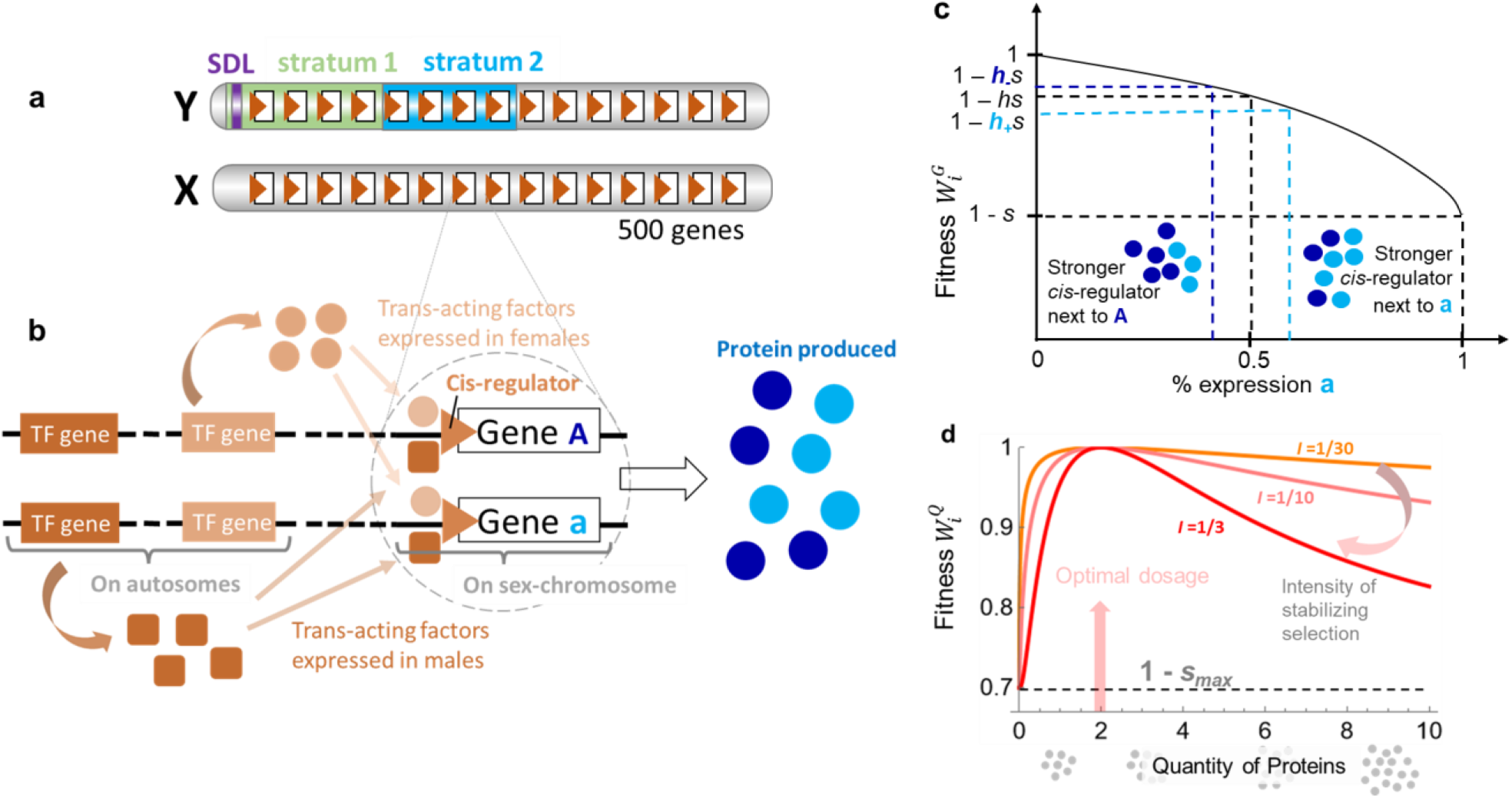
Graphical illustration of the model. (a) The model considers X and Y sex chromosomes with a sex determining locus (SDL) located at one end of the chromosome. The Y can evolve non-recombining strata (the figure illustrates a Y with two strata, in green and blue). 500 coding genes, each with its own cis-regulator are present on the Y and X. (b) The expression of each gene is symmetrically controlled by two autosomal trans-regulators, one expressed in males and the other in females, allowing for free evolution of dosage compensation. (c) Fitness effect of deleterious mutations: each allele of the coding gene expresses a given amount of protein determined by the trait values corresponding to its cis and trans-regulators (see Methods). The relative proportion of protein produced from the two alleles modulates the dominance level (illustrated by the *h_+_* and *h_-_* values on the y-axis) of the deleterious alleles that may occur in the coding sequence, as shown by the curve. (d) Fitness effect of expression level: The total amount of protein produced from a gene (as determined by trait values corresponding to its cis and trans-regulators) is under stabilizing selection (of intensity *I*) around an optimal dosage (set to 2), for each gene. Zero expression has a fitness effect of *s*_max_, corresponding to the maximal fitness effect of deleterious mutations that can accumulate in a gene (such that knocking out a gene by accumulating deleterious mutations has the same effect as reducing its expression to zero). This *s*_max_ value is set to 0.3 in all simulations. The contribution of gene *i* to fitness is given by 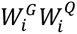, fitness effects are multiplicative among genes.

#### Regulatory theory for Haldane’s rule and the large X effect

Figure 2 shows the decrease in fitness in male and female F1 hybrids relative to the fitness of male and female offspring from within species crosses. HR is rapidly observed in all cases, with strong asymmetries between the fitnesses of male and female F1 hybrids. LX is also predicted in all cases, since autosomes of equivalent size generate weaker incompatibilities (compare Fig 2A to Fig 2B-E). Genes with an effect limited to the heterogametic sex (i.e., involved in fertility rather than viability) also contribute to HR and LX, and in many cases, very strongly (Fig 2E, §2.4 in *27*). The case of sex-limited genes can also predict the occasional occurrence of Y-autosomes incompatibilities, as observed in some cases (§2.5 in *27*). In §1.5 in (*27*), we show that with larger population size and stronger intensity of stabilizing selection on dosage, the fitness asymmetry between male and female F1 offspring tends to be larger, but the overall decline in hybrid fitness is slower. In §1.6 and Fig S6 in (*27*), we show that asymmetries between reciprocal crosses often occur, as observed in “Darwin’s corollary” to HR (*8*). In this model, autosomes also contribute to hybrid breakdown, but less strongly and without generating an asymmetry between the homo and heterogametic sex (Fig 2A, §2.1 in *27*). This outcome results from the stochastic coevolution of cis- and trans-regulators of gene expression, and their divergence among species (Fig S2), which has repeatedly been emphasized as a mechanism generating hybrid incompatibilities (*38–50*).

**Fig. 2.**
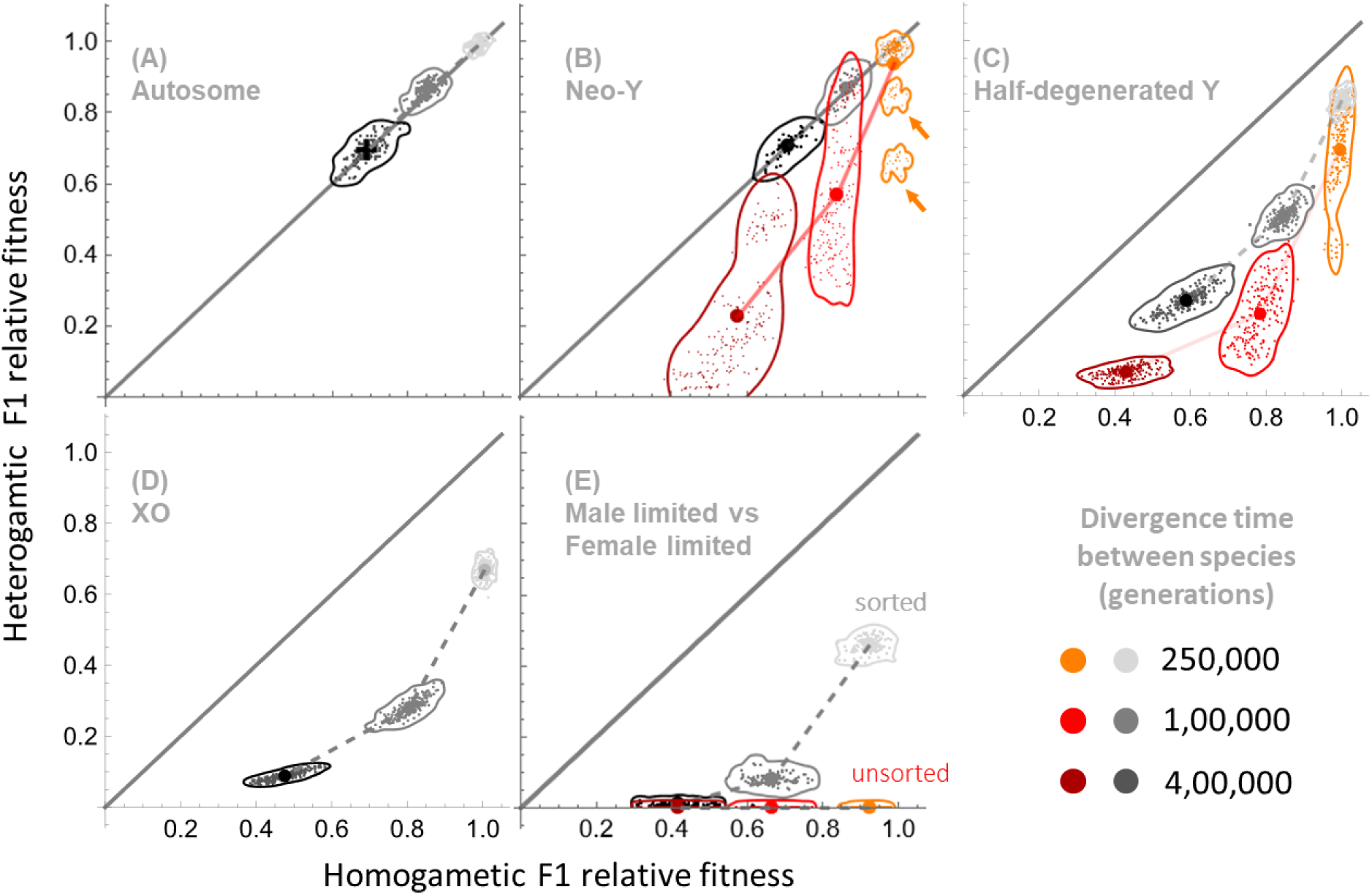
Hybrid fitness in crosses between Independently evolving species. The figure shows the fitness of homogametic (x-axis) and heterogametic F1 (y-axis), between species that have evolved independently for 2.5×10^5^, 10^6^ or 4×10^6^ generations (light, medium, and dark color on each panel, respectively). The data are obtained in each case using 15 independently evolving species and the 105 possible hybrid crosses (averaged in the two directions FxM and MxF for each of them). The fitness of hybrids is computed relative to the average fitness of intraspecific crosses for male and female offspring, respectively. Dots (or crosses for panel A) are mean values for all replicates, and small dots individual values. Contours represent the areas containing the individual values (the envelope is computed with a smoothed Gaussian kernel to facilitate vizualization). The x=y line is added to show equal fitness reduction in male and female F1. HR corresponds to cases where points fall below this line (the fitness of the heterogametic sex is lower than that of the homogametic sex in F1 hybrids). LX corresponds to a larger effect in panels B-E than the one shown on panel A for one autosome of equal size. (**A**) Autosomal case (see Methods). (**B**) Neo-Y case (see Methods). In gray, the simulations where inversions do not occur, so that the XY remains fully recombining throughout. In orange/red/darker red, inversions (and reversions) occur and recombination progressively stops between the X and the Y in each species independently (Fig S3 shows the progression of recombination arrest in the different replicates). After 2.5×10^5^ generations, only two species evolved a non-recombining stratum on the Y, and all hybrids involving those species had a lower fitness in the heterogametic sex (corresponding to two clouds of points, whose contours are indicated by the orange arrows). (**C**) Half degenerate case (see Methods). Note that regulatory divergence can accumulate initially for a larger number of dosage-compensated genes (initially 250), explaining why the fitness reduction on hybrids is larger for the same time period than in (B), where the simulations start with no dosage compensated gene. In gray, the simulations where inversions do not occur and where all the effect is therefore only driven by the shared ancestral stratum (the difference between the curves shows the contribution of newly evolving strata). (**D**) XO case (see Methods). In this case, there is no simulation with new inversions since there is no Y. Here too, regulatory divergence can accumulate initially for a larger number of dosage-compensated genes (500) than in (B) and (C), explaining the larger fitness reduction in hybrids. (**E**) Sex-limited case (see Methods). The figure shows the result of simulations with only female-limited genes (x-axis) or only male-limited genes (y-axis), starting from a fully non-recombining Y. These simulations are performed independently and are randomly paired between male-limited and female-limited simulations, for better comparison with the other cases (see Methods). In the “unsorted” case, each male-limited gene becomes independently degenerate and silenced on either the X or Y during the course of the simulation (see Fig S11), while in the “sorted” case, which gene is lost on the Y and which is lost on the X is initially decided, and the same for all replicate populations (see methods). The fitness of heterogametic F1 drops very quickly toward a value close to zero in the unsorted case.

**Fig 3.**
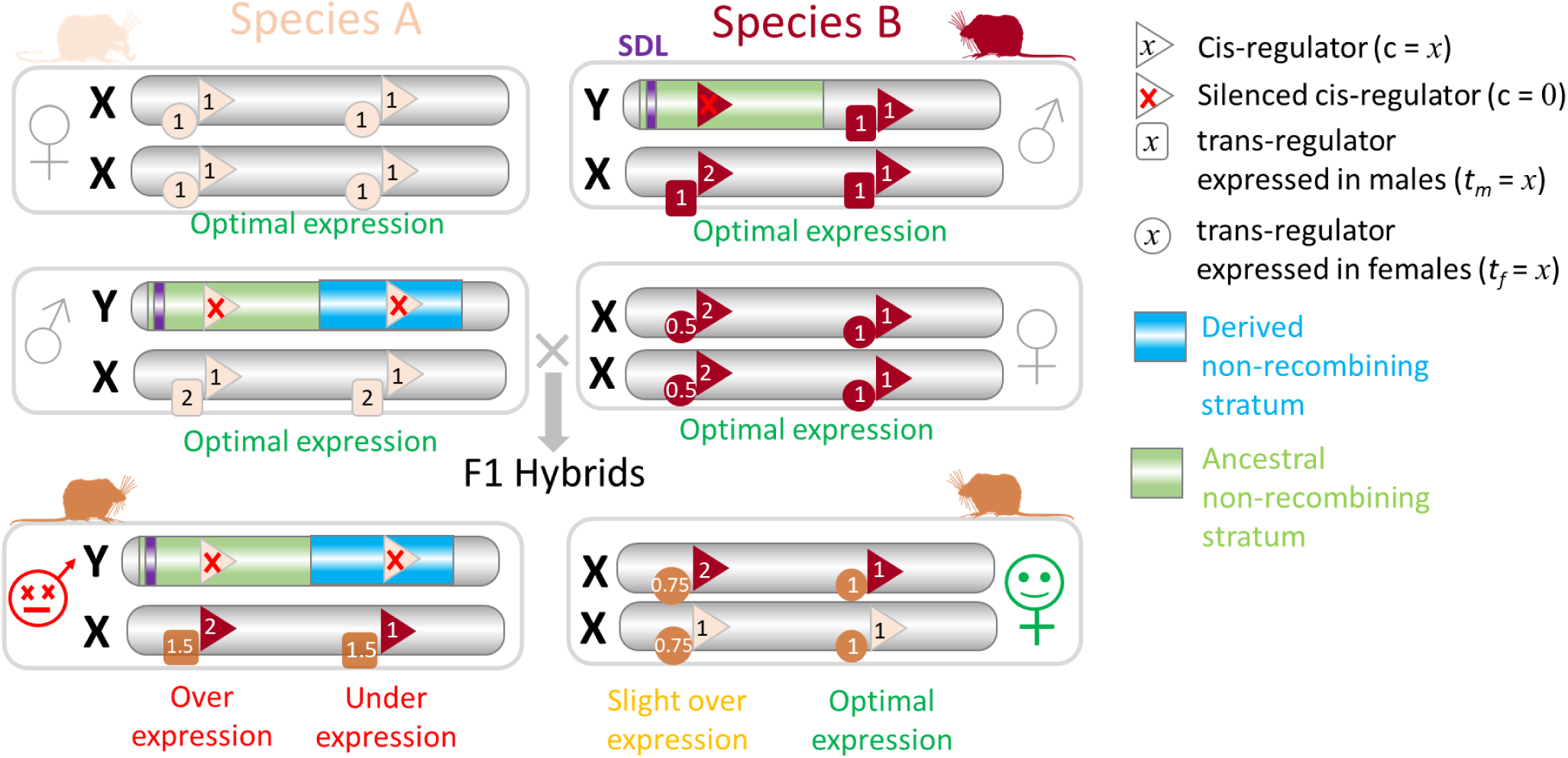
Misregulation of dosage compensated genes in F1 hybrids leads to Haldane’s rule and the large X effect. Cis and trans-regulators can stochastically diverge between two independently evolving species. This can occur throughout the genome and will reduce hybrid fitness in both males and female. However, regarding dosage-compensated genes, this regulatory divergence affects more the heterogametic than the homogametic F1 hybrid. Indeed, the former inherits, for a given dosage-compensated gene, a single cis-regulator (on the maternal X in XX/XY species), but two autosomal trans-acting factors (inherited from both parents), which creates an averaging mismatch between the two types of regulators, and misexpression of this X-linked gene. This is true for dosage-compensated genes present in the region of the X which stopped recombining with the Y in the ancestor of the two hybridizing species (green region on the figure) or in only one of them (blue region). Expression levels are computed as the sum of the cis-regulatory traits (numbers in the triangles) multiplied by the average of the (male or female expressed) trans-regulatory traits (numbers in squares and circles, respectively). Optimal expression is set at 2.

In this model, regulators on sex chromosomes evolve rapidly due to recombination arrest, chromosomal degeneration and the emergence of dosage compensation (*35*). As a consequence, differences in regulatory traits can rapidly evolve between species and cause dysfunctional regulation of dosage compensated genes in hybrids (Fig S2 shows the regulatory divergence observed in the different scenarios). But why would dysfunctional regulation disproportionately impact the heterogametic sex (HR) and cause a large X effect? For the latter, the answer is straightforward: if dysfunctional regulation of dosage compensated genes inordinately reduces hybrid fitness in the heterogametic sex, then the X chromosome, which is the only chromosome evolving DC, will necessarily have a disproportionate effect on hybrid fitness compared to autosomes (both in terms of the number and impact of genes involved in hybrid incompatibilities). Regarding HR, the disruption of the regulation of dosage compensated genes can be caused by the portion of the Y that is degenerate and compensated in only one of the two hybridizing species (Fig 2B, 2C) or by the portion of the Y that is degenerate and compensated in both species (Fig 2C, 2D), provided they exhibit some regulatory divergence. The effect per gene is on average larger in the first case (§1.4 in *27*), but the total effect depends on also on the number of genes in these different portions of the Y, so that both effects can contribute significantly to HR. We detail each case in turn.

#### Effect of a derived stratum present in a single species

First, consider the case where species *a* evolves a new non-recombining stratum on the Y, not present in species *b*. If a gene *g* is degenerate and compensated in species *a* but not in species *b*, then hybrid males will suffer from under-expression if they are 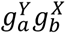, and overexpression if they are 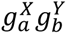, assuming codominant effects of the trans-acting factors. In the first case, these hybrids miss half the trans-acting factor required to fully achieve DC with a degenerated 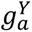 copy. In the second case, the hybrids inherit trans-regulators from species *a* increasing expression of gene *g*, despite having two functional copies of the gene. In all cases, the level of expression is closer to the optimum in females, regardless of the direction of the cross. This applies to the case where DC evolves by doubling the expression of gene *g* in males through a male-specific trans-acting factor (as with the *Drosophila* DC mechanism, Fig S7a) or where X chromosomes have doubled expression, one X being randomly silenced in females (as in the mammalian DC mechanism, Fig S7b). This argument applies more generally when evaluated quantitatively in a model encompassing these cases and their intermediates (§2 in *27*). The male/female fitness ratio is decreased proportionally to the number of genes in the stratum, the intensity of stabilizing selection on gene expression, and a constant that depends on the DC mechanism. The effect described here is similar to the verbal argument of Filatov (*51*), although clarifying the role of regulators.

#### Effect of an ancestral stratum present in both species

Consider now the case of two species that inherited from their common ancestor a portion of the Y that is non-recombining, degenerate, and dosage-compensated. In this case, misexpression of dosage compensated genes continues to cause a stronger fitness decrease in the heterogametic F1. In this scenario, the fitness decrease occurs when the regulatory traits, for a focal dosage compensated gene, have stochastically diverged between the two parental species. In F1 hybrids, these compensations will average out in females (who receive half the autosomal trans-acting factors and half the X cis-acting factors from each species) but not in males, who will have cis-acting factors on the X from a single species, but autosomal trans-acting factors from both species (Fig S8). Males will consistently under or overexpress more strongly compared to females. A quantitative model shows that the reduction in male/female F1 fitness is proportional to the number of genes on the stratum, the intensity of stabilizing selection on expression levels, and the between-species variance in the trait governing DC (§2 in *27*). This difference in averaging between sexes is similar to the mechanism generating HR in models of quantitative traits coded by multiple loci with additive effects (*25*, *26*). However, while these previous models may be seen as a particular case of the dominance theory (as they assumed a mutation on the X has the same effect when hemizygous in males and when homozygous in females), this is not the case in our model, since a mutation on a cis-regulator on the X does not interact with the same trans-regulators in males and females (§3.2 in *27*).

#### Sex-limited fertility genes

Sex-limited fertility genes can also contribute to the decrease in hybrid fitness of the heterogametic sex, with some interesting specificities. Female-limited genes on the X retain a standard diploid expression. The coevolution of their cis and trans regulators can slowly diverge between species, but this effect is relatively weak and comparable to what is observed on autosomes (§2 in *27*). The effect is much larger for male-limited genes, and even larger than for genes expressed in both sexes (Fig 2E). We also need to distinguish between cases where male-limited genes are present in a recently derived non-recombining portion of the Y in one species, or in an ancestral stratum. In the first case, gene expression divergence between X and Y-linked copies of genes present in the new stratum will generate a fitness cost for male hybrids in a similar way as for the non-sex-limited case (Fig S7a). However, a difference with the non-sex-limited case is that either the X or the Y-linked copy may be silenced following recombination arrest (while extinction of the X copy could not occur for genes expressed and required in both sexes). In the second case, male-limited genes may already have been silenced (either on the X or Y chromosome) after recombination arrest and before the split between the two species, in which case the divergence of their regulatory traits between species will generate a fitness cost for F1 males (Fig S9a). However, genes retaining diploid expression at the time of the split may strongly reduce the fitness of F1 males. Indeed, if the X-linked copy is subsequently silenced in one species while the Y-linked copy is subsequently silenced in the other, F1 males will either show a complete lack of expression or major overexpression of that gene (Fig S9b). A single essential gene in this situation could therefore cause male sterility in the heterogametic F1.

### Comparison to observations and other theories

Hence, HR, its “Darwin’s corollary” for reciprocal crosses (§2.6 in *27*), and LX could be caused by misregulation of sex-linked gene expression in hybrids (Fig 2, 3). Although the idea of DC disruption has been repeatedly discussed (*1*, *3*, *6*, *19*, *51–55*) and is close to the initial suggestion of Muller (*56*), it has never been quantitatively formalized, and usually considers that the DC mechanism is deficient in hybrids, not, more precisely, that the expression of dosage-compensated gene is deficient in hybrids. While most authors have been cautious about completely rejecting the DC disruption hypothesis, it has been usually dismissed as an important explanation (*5*, *7*, *13*) based on various arguments that we list and critically evaluate in §3.3 in (*27*). Most of the 13 criticisms we list seem either questionable or outdated (for instance, the idea that ZZ/ZW systems do not have DC while exhibiting HR). In most cases, they focus only on the direct disruption of the DC pathway, and not on the disruption of expression of dosage-compensated genes. Two arguments in this list remain relatively strong and require scrutiny.

#### Global versus local dosage compensation

The first argument states that in species with an ancestral ‘global’ (i.e., chromosome-wide) DC mechanism, there may be little scope for divergence in DC, for any focal gene, in recently diverged species. This would make a theory based on diverging DC less effective, especially in pairs of species where the same portion of the Y is non-recombining and where the cis-regulators involved in global DC do not seem to have diverged. However, DC disruption for a focal gene may still occur if target cis-regulatory sequences do not change but move location on the X (in systems like *Drosophila* with high-affinity sites ‘HAS’ targeted by the Male-Specific Lethal complex (*57*)) or if some genes escape the global DC mechanism. Current evidence indicates that chromosome-wide DC is the exception, not the norm, and even in these cases, many genes may escape global DC (*58–61*). As our theory shows, these genes should be scrutinized for their role in decreased hybrid fitness. Furthermore, global DC is often absent in the germline, where some DC nevertheless occurs for some genes (*62–66*). This is observed independently of the mechanism of meiotic sex chromosome inactivation (MSCI), which might have specifically evolved to control sex-chromosome meiotic drive during early meiosis (*67*, *68*), and the disruption of which has also been suggested to contribute to HR (*10*, *69–71*). This lack of a global DC mechanism for germline-limited genes in the heterogametic sex could explain why male fertility genes are major contributors to HR (in particular, male X-linked genes in *Drosophila melanogaster* are compensated independently of the global system based on the Male-Specific Lethal complex (*72*)). Indeed, without being tied to a global chromosomal level DC system, their local regulation could more easily diverge between sister species. The fact that male-limited genes can degenerate on the X and remain on the Y could also be a potent factor preventing somatic DC from being “used” in the germline: upregulating the X is certainly not fitting compensation for Y-linked genes expressed in the male germline. This could account for the observation that different DC systems evolve in somatic vs. germinal tissues (*66*, *73*). This leads to the prediction that HR should most often be based on hybrid sterility in groups with global somatic DC, while being more often based on viability in groups lacking it. Indeed, regulatory divergence of viability genes expressed in the soma may be limited in the presence of a global somatic DC mechanism, while local DC of sex-limited fertility genes (expressed in the germline) may diverge more easily. By contrast, HR should be more often based on viability in species lacking global DC, due the fact that many more genes affect viability than fertility. This could explain why birds and butterflies lacking global somatic DC (*74*) show more cases of HR through inviability compared to mammals and Diptera (*75*), an observation that has otherwise never received any explanation. Hence, the present theory could account for HR in species with global somatic DC, and also explains the contrasted prevalence of HR on sterility vs inviability in different groups, based on whether they evolved a global or local DC mechanism.

#### Comparison to the faster male theory

The second argument against a regulatory theory for HR concerns species with recombining sex chromosomes. In species without hemizygosity, it has been shown that HR is weaker but present compared to species with a more degenerated Y. This is the case, for example, in *Aedes* compared to *Anopheles* mosquitoes (*17*). This example has been used to support the “faster male” theory and to rule out the idea that the dominance theory alone could explain HR (*17*). The same argument would also argue against the theory described here. However, as recent evidence points out, in *Ae*. *aegyptii* the sex “locus” is a 1.5 Mb region with 30 genes in a 100 Mb non-recombining region encompassing the centromere of chromosome 1 and showing some divergence between males and females (*76–78*). Hence, these species may not entirely lack hemizygosity. Like above, genes in this region should be scrutinized for their role in decreased hybrid fitness. The other argument favoring the “faster male” theory is the overrepresentation of male sterility in HR, which is not a pattern directly following the dominance theory (*16*, *28*). However, our results also show that fertility genes can play a disproportionate role on HR, without invoking a “faster male” effect.

#### Comparison to the dominance theory

Beyond the faster male theory, the most strongly supported explanation for HR is the dominance theory. In particular, support for the dominance theory comes from experiments on *Drosophila* involving unbalanced females with attached X chromosomes. However, our theory makes similar predictions regarding these crosses (§2.3 in *27*), meaning that they cannot discriminate between theories. How do both theories compare beyond this? In marsupials and placentals, dosage compensation works by inactivating one X (the paternal or a random one, respectively). This is a difficulty for the dominance theory for viability, as females are somatically hemizygous (like males) in these cases. This is less of a concern for sterility, if the responsible genes are expressed in the germline where the inactive X is reactivated in mammals (*79*). This is not a major concern in our theory, where, as explained above, HR may result from the divergence in the global DC mechanism or from the few genes escaping global DC, whose regulation can more easily diverge between species (including genes only expressed in the male germline), but where the majority of genes subject to the same global DC will not cause any fitness reduction in hybrids. In the dominance theory, recessive incompatibilities occurring on genes escaping global DC may also contribute to HR. However, if such genes are rare, this contribution would be negligible, as the probability that a gene is both involved in an incompatibility and escapes global DC should be small: the vast majority of incompatibilities will concern genes subject to global somatic DC, so the fitness reduction in hemizygous males and females with X inactivation will tend to be similarly large. A second point relates to the underlying mechanism. Contrary to the dominance theory, which simply poses that a fraction of genetic incompatibilities are only expressed when one of the underlying loci is homo- or hemizygous (without providing a biological mechanism that would generate this type of interaction), our theory is based on a biological model of cis and trans regulator evolution, with underlying additive traits. Furthermore, direct empirical evidence is accumulating showing early regulatory evolution on sex chromosomes (*80–83*) and a role of misregulation of dosage compensated genes in hybrid dysfunction. For instance, in *Drosophila,* the key elements for DC—the MSL complex and the MSL-binding sites on the X—are fast evolving (*53*, *54*, *84*) as are *cis* and *trans* regulators of X expression (*85*). Indeed, Y-degeneration and DC appear sufficiently rapid and species-specific in related species with young and gene-rich sex chromosomes for the regulatory theory to work (*51*). Evidence is also accumulating linking the misregulation of sex chromosomes to HR in various hybrids (*10*, *71*, *86–94*). Misregulation of gene expression in hybrids has already been extensively studied (*38–50*, *86–88*, *91*, *94*, *95*), and improving the precision of techniques to measure gene expression will be crucial to critically evaluate our theory, and determine whether F1 hybrid fitness is indeed reduced by the misregulation of dosage compensated genes.

## Conclusion

Overall, and perhaps more importantly, HR is traditionally viewed as a composite phenomenon resulting from different causes (*1*, *3*, *5–7*, *13–16*, *16–19*), possibly distinct from the causes of the large X effect (*10*, *18*). As Coyne once wrote about HR and LX, “biology is not physics” (*14*), emphasizing that given the complexity of biological systems, unifying theories are unlikely. This conclusion may be true, but we suggest that it should perhaps be re-evaluated regarding HR and LX given the explanatory power of the regulatory theory presented here (Table 1). We show that a single model not only explains (1) recombination arrest on sex chromosomes, (2) degeneration of the Y, (3) the evolution of dosage compensation, as we previously showed (*35*, *36*), but also why, upon hybridization, (4) the heterogametic sex suffers more than the homogametic one, and (5) the X plays a disproportionate role in speciation. It may also explain why (6) HR is often asymmetrical between reciprocal crosses, (7) HR often involves fertility genes, (8) somatic and germline DC often differ, and why, (9) misregulation of gene expression on autosomes should follow HR in time and contribute (together with other types of post-zygotic isolation mechanisms) to complete reproductive isolation at a later stage of speciation. This theory therefore deserves to be further investigated empirically.

## Methods

Our baseline model (*96*) of sex chromosome evolution is the same as in ref. (*35*). In this study, we consider a population of 10,000 individuals (unless specified otherwise). Each individual has a pair of sex chromosomes with a sex-determining locus (located at one extreme end of the chromosome) and 500 genes that are subject to partially recessive deleterious mutations. These genes are regularly positioned along the chromosome (Fig 1a). The map distance between consecutive genes (*R_g_*) is set to 0.0005 (with 1 cM/Mb, this corresponds to a gene density of 20 genes/Mb, typical of plant and animal genomes). Recombination events are simulated by drawing the number of crossovers at meiosis from a Poisson distribution (whose mean is equal to the total map length) and drawing their position uniformly along the chromosome. Gene expression is regulated by cis-regulatory sequences (affecting expression only on the same chromosome as themselves, and whose strength is measured by a quantitative trait noted *c*) interacting with unlinked autosomal trans regulators (affecting expression of gene copies on both homologs, and whose strength is measured by a quantitative trait noted *t*) (Fig 1b). The map distance between each *cis*-regulator and its gene (*R_c_*) is 0.00005. All these elements mutate: mutation adds a unbiased Gaussian deviate *N*(0,*σ* = 0.2) to regulatory traits *c* and *t* (at rates *U*_*t*_ = *U*_*c*_/2 = 10^−4^, negative trait values are counted as zero). Mutations on coding sequences (occurring at rate *U*_*g*_ = 2. 10^−4^) are deleterious with an effect drawn from an exponential distribution with mean *s̄* = 0.05. The exponential distribution is used for simplicity and to ensure the occurrence of many small effect mutations. The relatively large mean is used to account for the absence of a class of loss-of-function mutations. Diffusion theory indicates that *s̄* should affect the results mostly through the *Ns̄* product and therefore that similar results would be obtained with smaller *s̄* and larger population sizes, provided that time is scaled by population size (*36*). These deleterious mutations can accumulate additively on each gene, up to a maximal value *s_max_* = 0.3 representing the fitness effect of losing that gene altogether. We assume that the dominance level of a deleterious mutation occurring in a coding gene is modified by the relative expression of the two copies of that gene, determined by the *cis* regulators linked to these two copies (*35*, *36*). Naturally, a deleterious mutation occurring in a less expressed gene copy is assumed to be less harmful than one in a more highly expressed copy. Specifically, and arbitrarily denoting by subscripts 1 and 2 the two alleles at a given gene in an individual, we assume that the fraction of the protein expressed from allele “1” at a given locus is *ϕ* = *c*_1_⁄(*c*_1_ + *c*_2_). This ratio measures the degree of allele-specific expression. With *ϕ* = 1/2 (i.e. with equally strong cis-regulators on both homologs), alleles are co-expressed, while a departure from ½ indicates that one allele is relatively more expressed than the other. Supposing that allele “1” is deleterious, then its dominance is given by *h*_1_ = *ϕ*^−*ln*(*h*)/*ln*^(^2^) (see Fig 1c), where *h* is the parameter measuring the baseline dominance of the fitness effect of deleterious mutations in a heterozygote when both alleles are equally expressed (with *ϕ* = 1/2, we have *h*_1_ = *h*). We use *h* = 0.25, corresponding to the average value observed across species (*97*). Fitness effects combine multiplicatively across loci (no epistasis). In order to allow dosage compensation on a gene-by-gene basis while maintaining symmetry between males and females, we assume that each gene is controlled by one male- and one female-expressed trans regulators, denoted *t_m_* and *t_f_* (the model does not address the question of evolving a global DC mechanism). We assume that the overall expression level (denoted *Q*) of each gene is subject to stabilizing selection around an optimal level *Q_opt_* = 2 with intensity *I* (see Eq. 4 in (*27*) and Fig 1d; we use *I* = 0.1 unless specified otherwise). For proper scaling, we ensure that, irrespectively of *I*, evolving zero expression has the same fitness effect as losing the gene altogether (i.e. at *Q* = 0, the fitness reduction is *s_max_*), which results in an asymmetry in the function describing this stabilizing selection (Eq. 4 in (*27*)). The expression level is given by *Q* = *t̄*_*m*_(*c*_1_ + *c*_2_) in males (where *t̄*_*m*_ is the average of the two alleles at the male-expressed trans regulator controlling the gene), and *t̄*_*f*_(*c*_1_ + *c*_2_) in females. Initially all *c* and *t* values are set to 1 (except when a portion of the Y is initially non-recombining, as explained below), leading to *Q* = 2.

We also introduce mutations suppressing recombination on a segment of the Y chromosome. For simplicity, these mutations are referred to as inversions, although they could correspond to other mechanisms causing recombination arrest. Inversions of any size are possible (their size is drawn from a uniform distribution over the recombining part of the Y), but we follow only those on the Y that include the sex-determining locus, which will necessarily be confined to males and cause recombination arrest. We assume that these inversions can accumulate, so that new inversions can occur on chromosomes carrying a previous inversion and thus extend the non-recombining region of the Y. Finally, we assume that reversions restoring recombination can occur, and for simplicity, that such reversions cancel only the most recent inversion (*35*). We consider that inversions occur at a rate *U_inv_* = 10^−5^, and that reversions occur 10 times less frequently. Table S1 recapitulates all parameters used and their baseline values.

We consider three main scenarios. In the first (“neo-Y”), the sex chromosome pair has just acquired the sex-determination locus. We follow the independent evolution of this recent Y in different replicate species (we call them species for simplicity, but these are, at least initially, only independently evolving populations). After a given number of generations, F1 hybrids are generated between these independently evolved species, in the two directions of the cross (M x F and F x M), and their fitness is compared to that of male and female offspring produced within species. The second scenario (“half-degenerate case”) corresponds to the case of a partially degenerated Y chromosome: it is equivalent to the first scenario, except that 50% of the Y is already non-recombining, fully silenced (*c*_Y_ = 0) and fully dosage compensated in the common ancestor of the diverging species (with Drosophila-like DC, i.e. with the trans-acting factors expressed in males set to *t_m_* = 2 and no change in females *t_f_* = 1). Finally, the third scenario (“XO”) assumes that 100% of the Y is already non-recombining, fully silenced and fully dosage compensated in the ancestor (this is equivalent to considering diverging species with a XX / XO sex determination system); in this case we also use *t_m_* = 2, *t_f_* = 1 as the initial condition). These scenarios allow us to compare hybrid fitness between species at different stages of sex chromosome evolution.

We also investigate the case of genes with male- or female-limited effects to represent genes involved in fertility rather than viability. In this case, we suppose that the sex chromosomes are initially not recombining. We performed simulations with male and female-limited genes independently. When a simulation runs with only male-limited genes, the fitness of females stays at 1 throughout, and reciprocally for female-limited genes. For the comparison, we therefore randomly paired male-limited and female-limited simulations. For female-limited genes, all genes are initially non-degenerate and with fair diploid expression. Degeneration does not occur on the X. Whether degeneration occurs on the Y is irrelevant since the genes are all female-limited and therefore not expressed in males. For male-limited genes (assuming a XY system, the reverse situation with female-limited case would occur for a ZW system), we considered two initial conditions: (1) all genes are initially non-degenerate and with fair diploid expression. During the course of these simulations, each male-limited gene will degenerate and become silenced on either the X or the Y, independently in different populations. We term this initial condition the “unsorted” case since it is initially not decided which gene will be lost on the Y or on the X. Results show that c.a. 90% of genes end up being degenerate on the Y (and 10% on the X, Fig S11). This asymmetry is due to selective interference on the non-recombining Y facilitating the initial accumulation of deleterious mutations on Y copies, which may be amplified by the fact that the effective size of the Y is three times smaller than the effective size of the X. In the second initial condition, we suppose that 50% of the genes are degenerate on the Y (and dosage compensated on the X with *t_m_* = 2) while 50% are degenerate on the X (and dosage compensated on the Y with *t_m_* = 2). We term this initial condition the “sorted case”, meaning that it is already decided which genes are degenerate on the X or Y (and it is the same for all replicated species).

In addition to these scenarios, we performed simulations with a fully recombining autosome instead of a sex chromosome pair (without including new inversions on this autosome but including the possibility of regulatory evolution, with one male and one female unlinked trans-acting factor per gene). Note that in these simulations, the sex of each newly formed individual is drawn randomly (with an expected 1:1 sex ratio). Another set of simulations involves sex-chromosomes, but without considering any new inversion (in the XO scenario, this case is irrelevant since the whole Y is already non-recombining). Finally, we also investigated the effect of varying population size and the intensity of stabilizing selection on dosage (§1.5 in *27*).

## Acknowledgments

We thank D Presgraves, C. Haag, M. Kirkpatrick, S. Otto, M Raymond, J Welch and anonymous reviewers for discussions and/or comments, K. McKean for editing and L Meslin for illustration. We thank the CNRS ABiMs cluster.

## Author contributions

Original ideas TL, DR; Model conception TL, DR; Code DR, TL; Simulations TL; Data analyses TL; Interpretation TL, DR; First draft, editing and revisions TL, DR; Project management and funding TL.

## Competing interests

the authors declare no conflict of interest.

### Data and materials availability

(*96*).

## Supplementary material

### 1 Additional methods and results

#### 1.1 Individual fitness and parameter values

The fitness of individuals is the product of two components: one resulting from the presence of deleterious mutations in genes 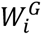 and one resulting from the expression level of that gene 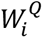. For a gene *i*, the contribution of deleterious mutations is

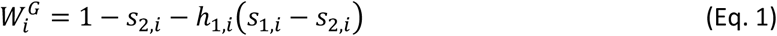

where, by convention, subscript 1 denotes the most deleterious allele of the two present in a given individual for that gene *i*. The dominance coefficient of this most deleterious allele depends on the relative expression level of both alleles. Noting the fraction of the protein expressed from allele 1

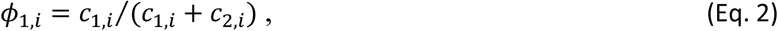

where *c*_1,*i*_ and *c*_2,*i*_ are the cis-regulator strengths associated to the two alleles, this dominance is given by

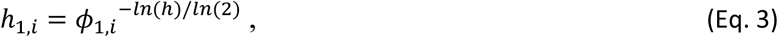

where *h* is a parameter measuring the baseline dominance of the fitness effect of deleterious mutations in a heterozygote when both alleles are equally expressed.

For a gene *i*, the contribution of the expression level of gene *i* to fitnesss is given by

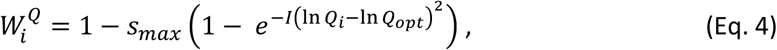

where the total expression level *Q*_*i*_ for a gene *i* is equal to (*c*_*X*,*i*_ + *c*_*Y*,*i*_)*t̄*_*m*,*i*_, where *t̄*_*m*,*i*_is the average strength of the *trans*-regulators expressed in males, while *c*_*X*,*i*_ and *c*_*Y*,*i*_ are the strength of cis regulators associated with the X and Y-linked copies of gene *i*, respectively. Symmetrically, it is (*c*_*X*1,*i*_ + *c*_*X*2,*i*_)*t̄*_*f*,*i*_ in females. *I* measures the intensity of stabilizing selection on the expression level, and *s_max_* is the fitness effect of completely disrupting or silencing the gene (see Methods).

The same equations are used to measure hybrid fitness. Hybrids are made by sampling X and Y chromosomes from the simulation at a given time, paired with chromosomes sampled from another replicate simulation (at the same time) to produce male and female F1 hybrids. Parameter values used in the simulations are given in Table S1.

**Table S1.** Parameter values. Model’s parameters, their notation, description and values (in bold their default value when several values were considered).

| Parameter | Description | Value |
| --- | --- | --- |
| $N_g$ | Number of genes | 500 |
| $N$ | Number of individuals | **10000**, 5000, 20000 |
| $\bar{s}$ | Average fitness effect of deleterious mutations in coding genes | 0.05 |
| $h$ | Baseline dominance of deleterious mutations in coding genes | 0.25 |
| $s_{max}$ | Maximum cumulated effect of deleterious mutations in a given coding gene | 0.3 |
| $I$ | Intensity of stabilizing selection on dosage | **1/10**, 1/30, 1/3 |
| $\sigma$ | Standard deviation of the phenotypic effect of regulatory mutations | 0.2 |
| $Q_{opt}$ | Optimal expression level for coding genes | 2 |
| $U_g$ | Mutation rate per coding gene per generation | $2 \cdot 10^{-4}$ |
| $U_c$ | Mutation rate per cis-regulator per generation | $2 \cdot 10^{-4}$ |
| $U_t$ | Mutation rate per trans-regulator per generation | $10^{-4}$ |
| $U_{inv}$ | Rate of inversion per chromosome per generation | $10^{-5}$ |
| $U_{rev}$ | Reversion rate of inversions per generation | $10^{-6}$ |
| $R_g$ | Map distance between consecutive coding genes | $5 \cdot 10^{-4}$ |
| $R_c$ | Map distance between a coding gene and its cis-regulator | $5 \cdot 10^{-5}$ |

#### 1.2 Regulatory divergence

The regulatory system for a given species and a given gene can be characterized by the vector **r** = (*t*_*m*_, *t*_*f*_, *c*_*X*_), where *t*_*m*_, *t*_*f*_, *c*_*X*_ are the trait values for the male trans-regulator, the female trans-regulator and the cis-regulator on the X. On the Y, the cis-regulatory trait tends towards zero after recombination suppression and thus does not contribute to regulatory variation among independently evolving species (before recombination suppression, *c*_*Y*_ equals *c*_*X*_ and therefore does not bring any additional information on regulatory divergence). At a given time and for a given gene, we can therefore measure the regulatory divergence among independently evolving species as the variance of the regulatory system among replicates. Specifically, we measured it as the mean over replicates *i* of (**r**_***i***_ − **r̄**). (**r**_***i***_ − **r̄**). This divergence can then be averaged across genes for a specific region of the sex chromosome. Fig S2 shows this measure of divergence every 40,000 generations, over windows of 100 consecutive genes on the sex chromosomes, for the simulations represented on Fig. 2.

On autosomes or in the recombining region of the Y, regulatory divergence occurs by the stochastic variation of regulators. Selection ensures that cis and trans regulators match to produce the required amount of protein (i.e. that the product of their strength is near the optimal value). However, their absolute value is relatively free to evolve. It is a typical case of evolution “freedom” where traits evolve along a “neutral ridge”, to take a fitness landscape metaphor (*98*). However, when recombination suppression evolves on a region of the Y, then cis-regulatory traits in that region evolve to zero (silencing) and X-linked cis-regulators and trans-regulators quickly evolve to maintain optimal expression in both sexes (evolution of dosage compensation, see Eq. 5). The evolution of dosage compensation is quickly perturbing the regulatory system, causing a temporary acceleration of regulatory divergence. Fig. S2 shows this outcome quantitatively. Regulatory divergence is accelerated on sex chromosomes compared to autosomes, although not dramatically so. This acceleration occurs because sex chromosomes have a lower effective population size than autosomes (see also Fig S5b for the effect of N on regulatory divergence). This divergence is also temporarily accelerated when the regulatory system is perturbed by the evolution of dosage compensation on new non-recombining strata.

**Fig. S2.**
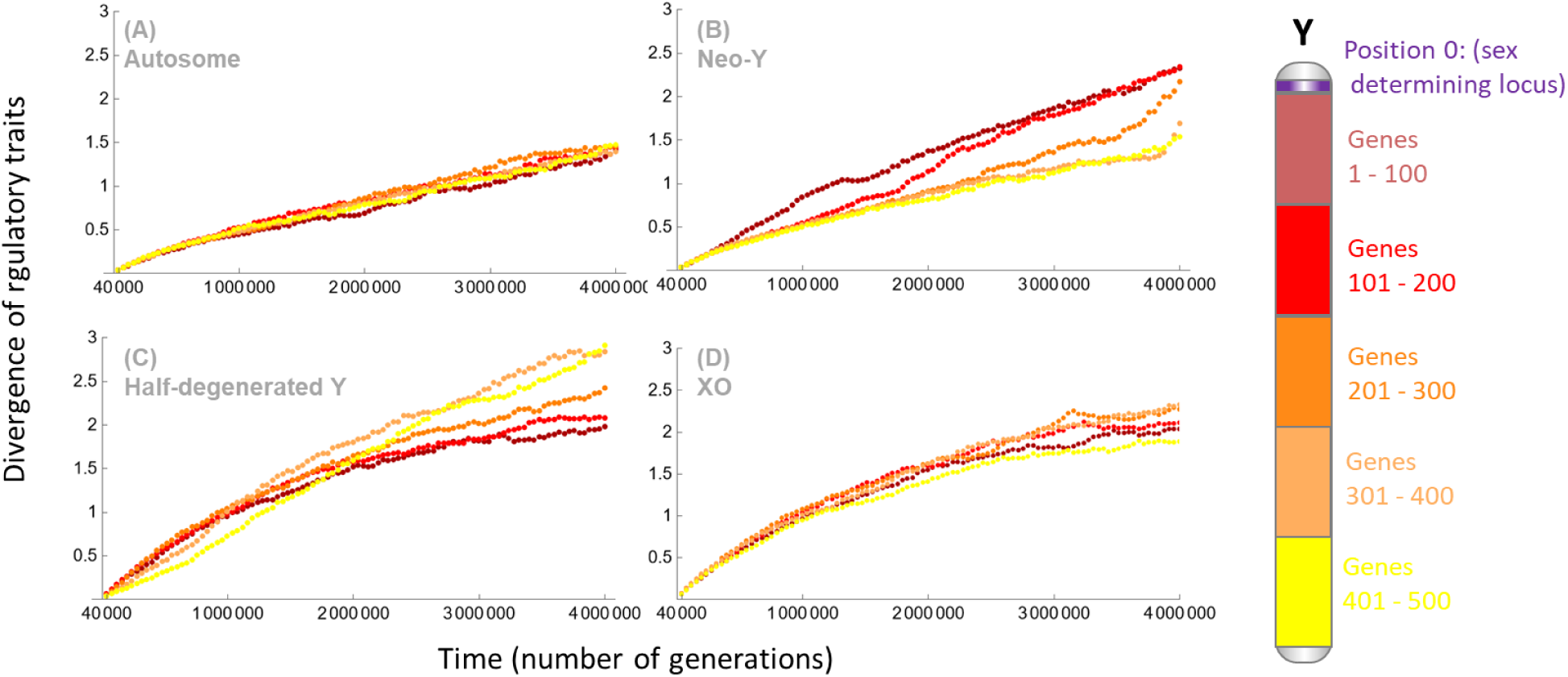
Divergence of regulatory traits through time. The x-axis is time (number of generations). The y-axis is a measure of the divergence of regulatory traits through time in independently evolving species. To compute it, the chromosome is divided into five equally sized regions: region 1 (brown) includes genes 1 to 100, region 2 (red) genes 101 to 200, region 3 (orange) genes 201 to 300, region 4 (light orange) genes 301 to 400, region 5 (yellow) genes 401 to 500. The divergence measure is given every 40 000 generations averaged over the 100 genes in these five regions, from the same simulations than those reported on Fig. 2: (A) Simulations with an autosome, (B) starting with a neo-Y, (C) starting with a half-degenerate Y, (D) with a fully degenerate Y. When recombination suppression evolves in some replicates, but not all, it causes a slight acceleration of regulatory divergence, as expected. This effect can be seen on simulations starting with a neo-Y (panel b). Recombination suppression occurs first in some replicates in the first region (brown), then in the second (red) etc. It can also be seen on simulations starting with a half-degenerated Y (panel c). Recombination suppression occurs first in some replicates in half of the third region for genes 250 to 300 (orange), then in the fourth (light orange) and the fifth (yellow). Note that half the third region (genes 201-250) was already degenerate and dosage-compensated at the start of the simulation, and thus does not contribute to the acceleration of divergence caused by the extension of the non-recombining region. The fact that this curve is below the curve for genes in positions 301-400 results from this effect.

#### 1.3 Evolution of new non-recombining strata

In neo-Y (Fig 2B) and half-degenerated Y scenarios (Fig 2C), with new inversions and reversions, recombination suppression evolves independently in the different replicates. Some inversions fix in the population and are not reversed : they are “stabilized” (see ref. (*35*) for details about the underlying processes leading to these fixations and stabilisation of these new strata). The size of newly occuring inversions is drawn from a uniform distribution over the recombining fraction of the Y chromosome. However, the process of fixation and stabilization favors inversions with intermediate but relatively small size. The size of each new stabilized stratum can be followed independently in each replicate through time. This is illustrated on Fig S3.

**Fig S3.**
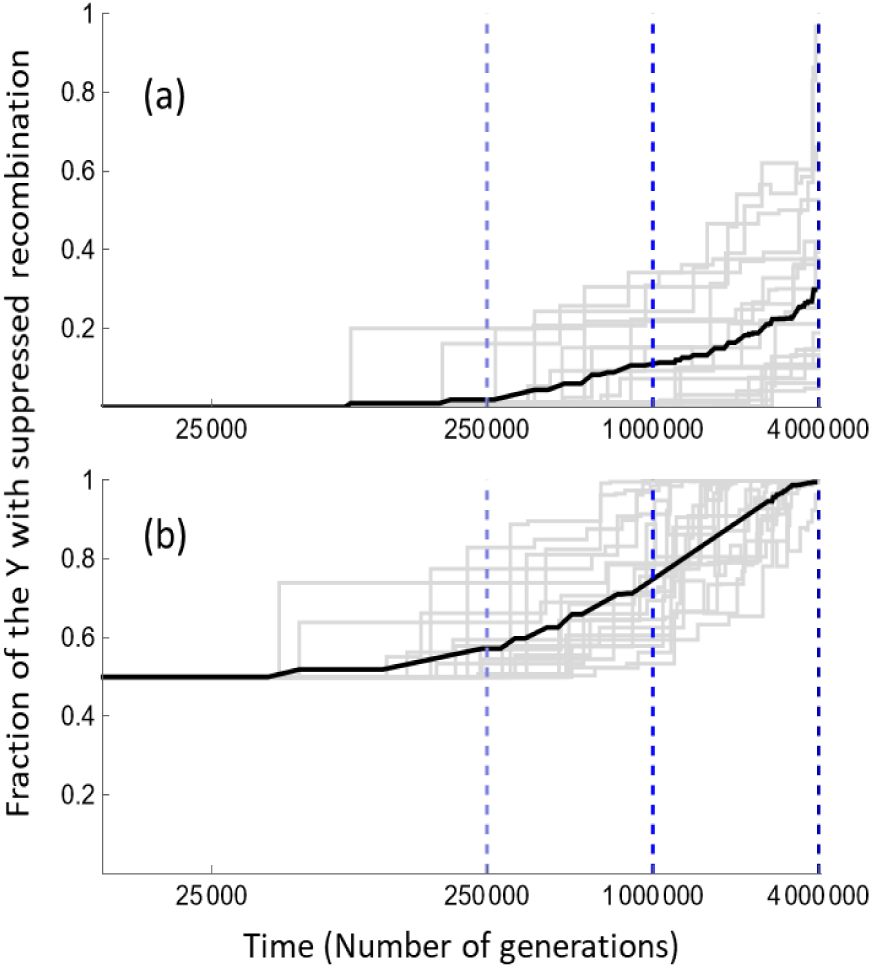
Evolution of recombination suppression in neo-Y and half-degenerate Y scenarios. **(a)** Evolution of recombination suppression in the neo-Y simulations corresponding to Fig 2B. The x-axis is time in number of generations, in log-scale, with the three time points illustrated on Fig 2 shown by the vertical dashed curves. The y-axis is the fraction of the Y with suppressed recombination corresponding to the accumulation of several strata. The gray lines represent individual replicates (F1 hybrids are generated by crosses among these replicates) and the black curve the mean across replicates. **(b)** Same as in (a), but for the simulations starting with a ‘half-degenerate’ Y, corresponding to Fig 2C.

#### 1.4 Fitness effect per gene in derived and ancestral strata

Consider two species evolving in a neo-Y scenario (their common ancestor has a fully recombining Y). Suppose that after *t* generations of divergence, species 1 evolves a non-recombining region up to position *z*_1_ on the Y chromosome, and species 2 up to *z*_2_. We can define the region up to Min(*z*_1_ *, z*_2_) as a “parallel” region. This region is non-recombining in both species but has evolved independently in each. The region between Min(*z*_1_ *, z*_2_) and Max(*z*_1_ *, z*_2_) can then be defined as a “private” region (i.e. this region is non-recombining in only one of the two species). These parallel and private regions are both caused by “derived” strata, i.e. strata occurring independently in the two species after speciation (Fig S4c). The mean effect of a gene (on the fitness of F1 individuals) from parallel and private regions can be computed by averaging over genes in parallel or private regions in all species pairs. Similarly, we can consider two species evolving in an XO scenario. Their common ancestor has a fully non-recombining, degenerate and dosage-compensated Y (with a given dosage compensation *t_m_*). After *t* generations of divergence, the mean effect of a gene (on the fitness of F1 individuals) within this “ancestral” region (of 500 genes) can be computed and averaged over all pairs of species. All these measures can be made independently for females and males (i.e. for the homo- and heterogametic sex, respectively).

Figure S4a shows these mean fitness effects per gene in parallel, private and ancestral non-recombining regions. The effect of genes is of the same order of magnitude among derived strata (private or parallel), with a large difference between their effects in male and female F1. The measures show some noise, especially initially when few derived strata have evolved (Fig S4b shows the average number of genes present in parallel and private strata over time). The effect of genes in the ancestral strata is also greater in males than in females, but the male effect is smaller (compared to the derived strata). This observation confirms equations 11-13 where the expected effect on ancestral strata per gene is expected to be weaker than on derived strata (see §2.2 below). Note that the mean effect per locus is small here. However, HR may not necessarily be caused by many small effects. Few dosage sensitive loci may contribute strongly to HR, and in some cases, a single gene could even cause sterility in hybrids (see §2.4 about the effect of sex-limited genes).

**Fig S4.**
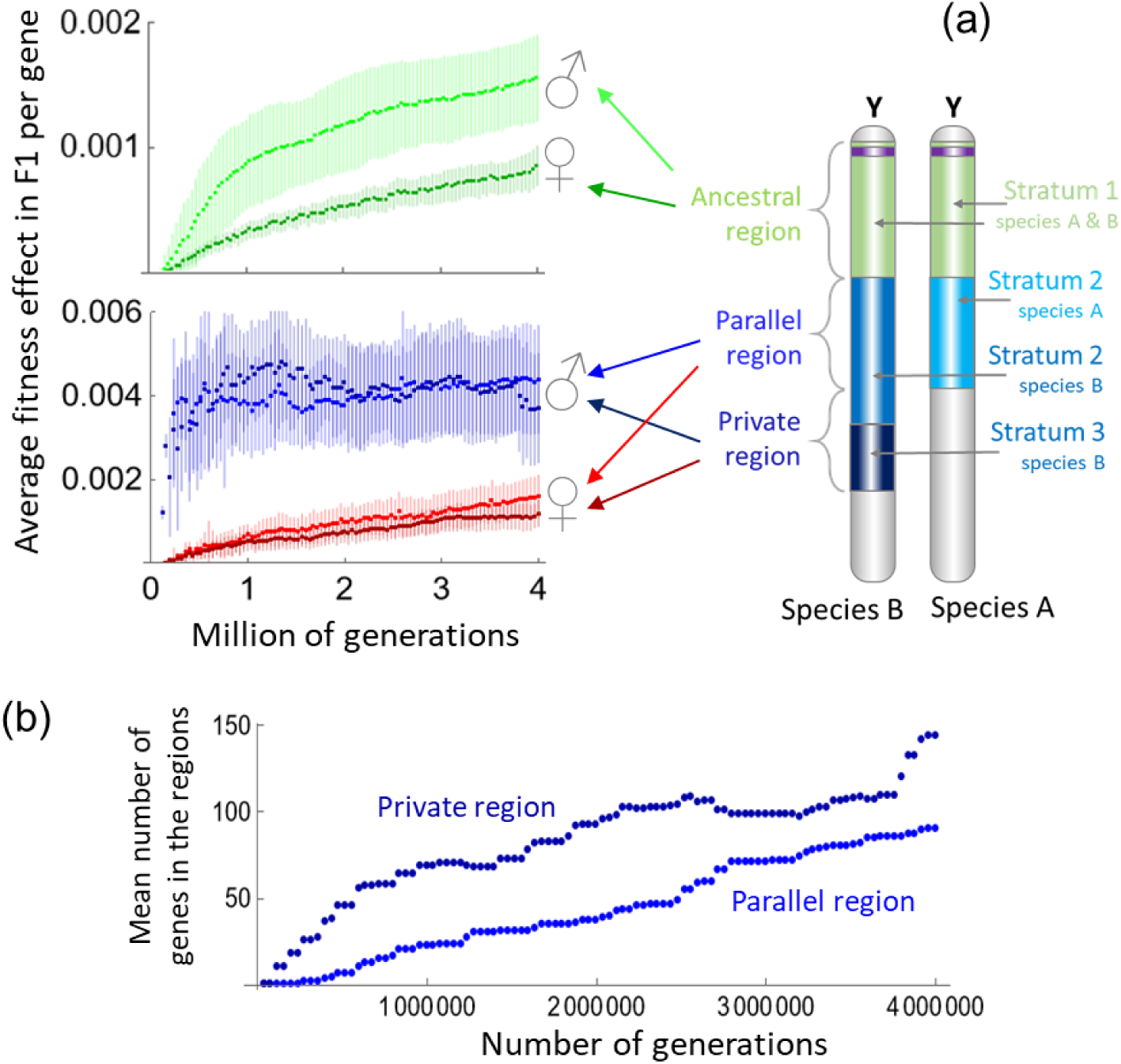
Effect per gene and number of genes in different non-recombining regions. (a) Mean effect of genes on F1 fitness in different non-recombining regions (parallel, private, ancestral, see their definition on the sketch of two Y chromosomes from two different species with different strata) after a given time of divergence between species (in number of generations, x-axis). This is computed in F1 XY males and F1 XX females (as shown by the male and female symbols). The mean is reported (dots) plus or minus the standard deviation of these per-gene values (across genes). Note that the scale of the y-axis differs for the ancestral region (top graph) and for the parallel/private regions (bottom graph). The data are from the neo-Y simulations shown on Fig 2B (for parallel and private region) and Fig 2D (for ancestral region). (b) Mean number of genes in parallel or private regions in neo-Y simulations.

#### 1.5 Effect of population size and intensity of stabilizing selection on hybrid fitness

We investigated the effect of varying population size *N* and the intensity of stabilizing selection *I* on dosage on the fitness of F1 hybrids. In the main scenarios reported on Fig. 2, population size was set to 10,000 individuals and *I* = 1/10. We performed additional simulations with 5,000 or 20,000 individuals in scenario 1 (neo-Y) and 3 (XO) to investigate the effect of increasing or decreasing population size. We also performed simulations with *I* = 1/30 or 1/3 in scenario 1 (neo-Y) and 3 (XO) to investigate the effect of increasing or decreasing the intensity of stabilizing selection (Fig S5a). Other parameters have predictable effect on the outcome and were not investigated. Notably, increasing the number of loci or the effect of a gene knock-out (*s*_max_) will directly increase the total fitness effect seen in hybrids (see Eq. 11) and varying mutations rates will change the timescale. The effect of the parameters on the process of Y evolution was also investigated in previous studies, and will influence the process in a predictable way in the scenarios involving the formation of new sex-chromosome strata (*35–37*). The effect of varying *I* or *N* is less straightforward as it can be expected to have opposing effects on the outcome. Indeed, increasing both *N* and *I* speeds up the formation of new strata (*35*). This is due to the fact that dosage compensation emerges more quickly on new strata in these cases, which protects them against reversion and the reestablishment of recombination (*35*). However large *N* and *I* may also subsequently limit regulatory divergence by cis-trans coevolution, since this coevolution involves crossing small “fitness valleys”, which is easier with weak selection on dosage and smaller population size. Fig S5b shows that this trade-off does indeed occur: larger *N* and *I* lead to faster evolution of recombination suppression in the neo-Y scenario, but reduce the speed of regulatory divergence. In the XO simulations, only the later effect is involved (the situation is equivalent to have a fully degenerate and non-recombining Y, there are no new strata). We therefore observe a large asymmetry between the homo and heterogametic sex, and their fitness decreases when regulatory divergence is faster (i.e. for low *N* and low *I*). In the neo-Y simulations, the regulatory divergence is also faster for low *N* and low *I*, however, the rate of acquisition of new strata is also reduced in these cases, so that the asymmetry between the homo and heterogametic sex is less pronounced for low *N* or low *I*.

**Fig S5a.**
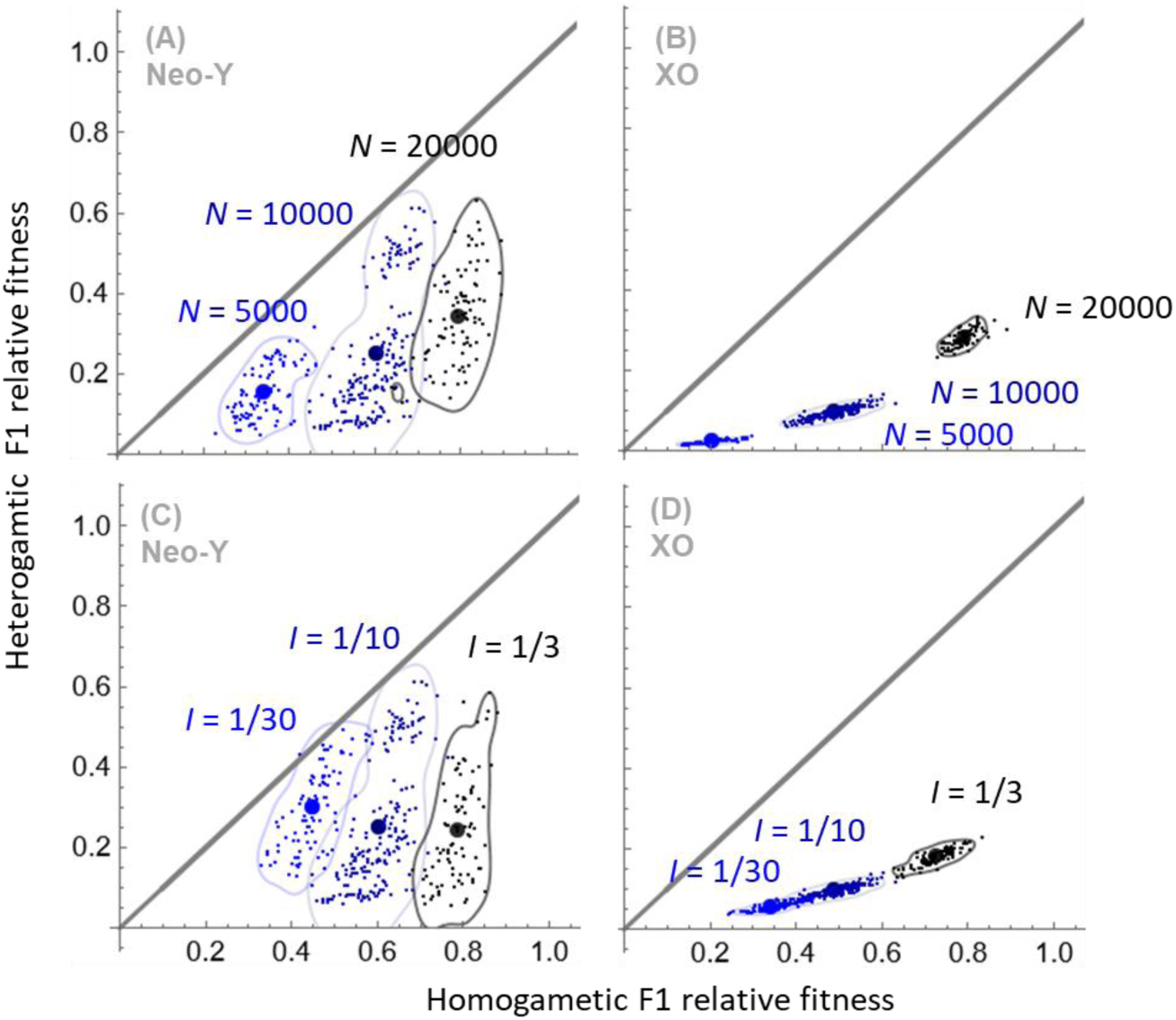
Hybrid fitness in crosses between Independently evolving species for varying population size *N* and intensity of stabilizing selection on dosage *I*. The figure is built like Fig 2 in the main text and shows the relative fitnesses of homo- and heterogametic F1 hybrids in neo-Y (panels A and C) and XO scenarios (panels B and D) after 4 million generations. Panels A and B show results for varying population sizes (values indicated on the plots, with *I* = 1/10). Panels C and D show results for varying intensity of stabilizing selection (values indicated on the plots, with *N* = 10,000). Each dot represents F1 hybrids from a specific species pair. Larger dots represent mean values and contours around dots (computed with a smoothed Gaussian kernel) are added to facilitate visualization.

**Fig S5b.**
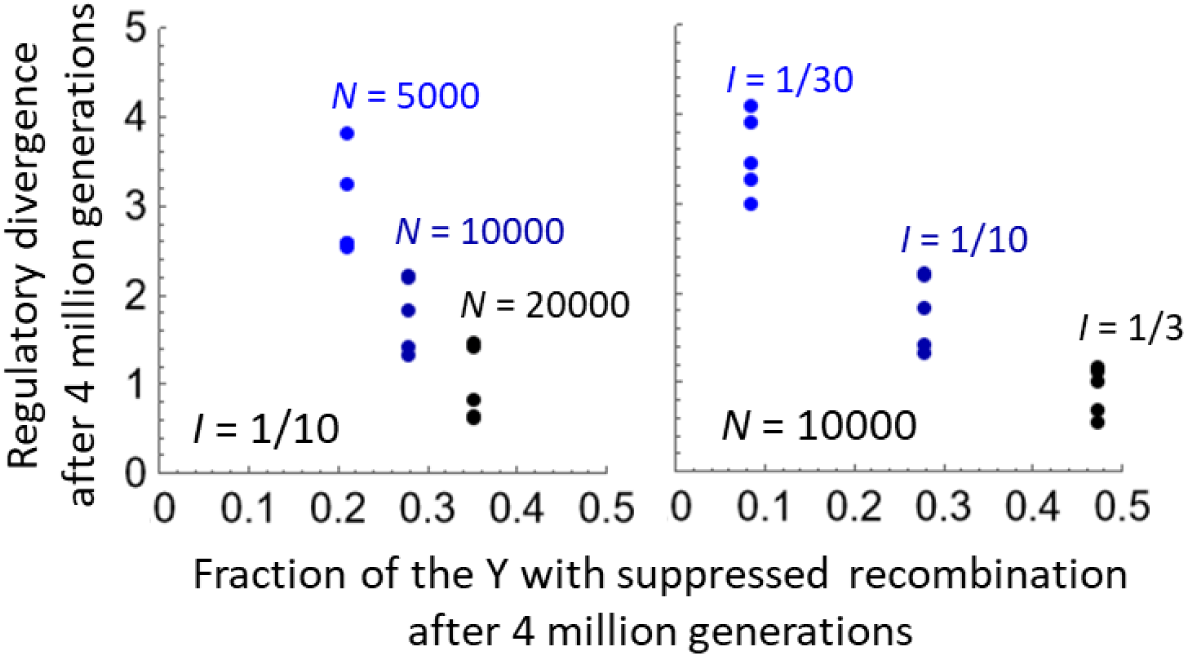
Trade-off between the rate of regulatory divergence and the rate of accumulation of new non-recombining strata. The x-axis shows the mean fraction of the Y that became non-recombining in the neo-Y simulations, after 4 million generations, for simulations with varying population size (*N*), but constant intensity of stabilizing selection on dosage (*I*), left panel, or the reverse, right panel (see parameter values on the figure). The y-axis shows the regulatory divergence, computed as on Fig S2, after 4 million generations, for the same set of simulations. The five dots correspond to the five 100-genes regions on the chromosome, as defined on Fig S2. The region closer to the sex determining locus tend to show larger regulatory divergence.

#### 1.6 Darwin’s corollary

The fitness of F1 hybrids often differs significantly between reciprocal crosses. For instance, Turelli and Orr (*99*) estimated that 15% of cases of Haldane’s rule in *Drosophila* involved a strong asymmetry (the male F1 being sterile or inviable in only one direction of the cross between the two hybridizing species). A similar pattern is seen in *Anopheles* (*17*) and could be even more prevalent in *Lepidoptera* (*100*) or *Silene* (*8*). Turelli and Moyle termed this pattern “Darwin’s corollary”, complementing Haldane’s rule and the large X effect in the rules of speciation (*8*). In our model, fitness asymmetries between reciprocal crosses often arise. The general reason for this asymmetry is that the heterogametic F1 will typically suffer from over-expression in one direction of the cross and from under-expression in the other. This pattern occurs in all cases (see detailed examples in Fig S7-S9). Everything being equal, under-expression is more deleterious than overexpression in our model, as is likely in most biologically plausible situations. In the extreme case, e.g. when a male-limited gene is entirely missing in one direction of the cross, male fitness is much more reduced than when its expression is doubled in the other direction of the cross (Fig S9b). The asymmetry in reciprocal crosses is particularly strong when a discrete event of large effect occurs in one species but not in the other (similarly, Muller noted that asymmetry between reciprocal hybrids must involve loci of large effect (*101*)). This occurs for instance when a species acquires a new non-recombining stratum and not the other. In this case, the F1 male inheriting the more degenerate Y will suffer more than the F1 male in the reciprocal cross (Fig S7a, S7b). This effect will occur mostly at intermediate steps of Y chromosome degeneration, i.e. when species have different strata, and before F1 male fitness is not too strongly reduced in all crosses (Fig S6a). Another case particularly conducive to strong fitness asymmetries in reciprocal crosses is when a male-limited gene is missing in one direction, as explained above. However, if there are many male-limited genes, then different genes will be lost on the X or Y in the different species, and the fitness of all F1 males will be reduced. Darwin’s corollary should be strongest when the number of male-limited genes on sex chromosomes is not too large, maximizing the sampling variance of those genes and therefore the fitness asymmetries in the reciprocal crosses (compare Fig S6b(a) and S6b(c), with 500 and 50 male-limited genes on the chromosome, respectively). When the fitness reduction in F1 males results from many small effects, fitness asymmetries in reciprocal crosses tend to be weaker. This is the case for instance when regulatory divergence occurs on a chromosome that is already fully dosage-compensated (see Fig S6b for XO simulations). Overall, our model predicts substantial fitness asymmetries in reciprocal crosses, and at intermediate times of species divergence, which is in line with the available observations.

**Fig S6a.**
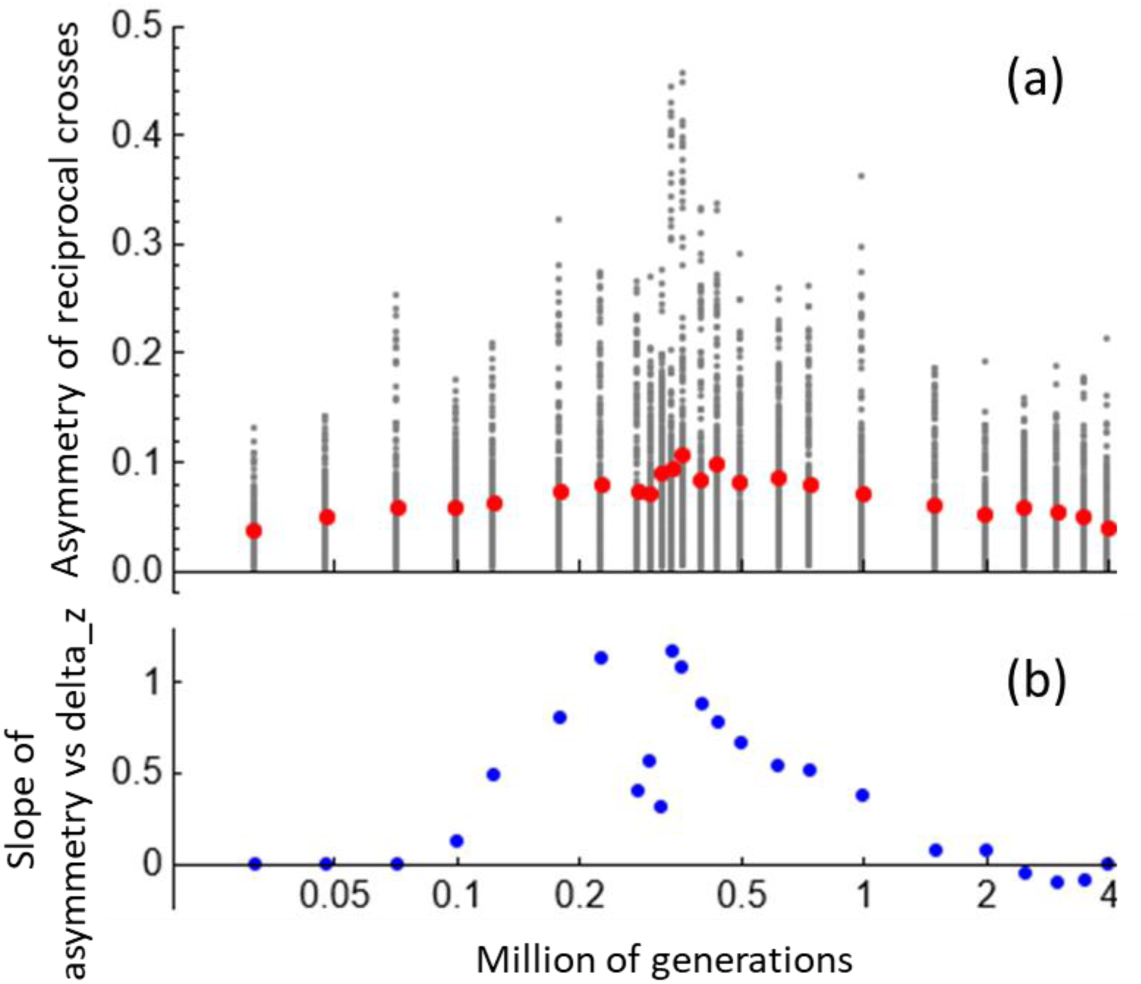
Darwin’s corollary in the neo-Y scenario. (**a**) Crosses asymmetries in neo-Y simulations (same simulations as the one illustrated in Fig 2B). The axes show the absolute value of the difference in the fitness of males produced in reciprocal crosses (y-axis) against time (x-axis, in millions of generations, in log-scale). A larger value means that the male fitness reduction is larger in one direction of the cross. Gray dots are the individual values obtained from all hybrids among independently evolving populations; red dots indicate mean values. (**b**) The graph shows the slope of the regression of the (signed) fitness difference between males produced in reciprocal crosses against the (signed) difference in the size of the non-recombining region (*z*) between the Ys in the two populations (y-axis). A positive slope indicates that males suffer more in the direction of the cross where the father carries a Y with a larger non-recombing region (i.e. a larger z value).

**Fig S6b.**
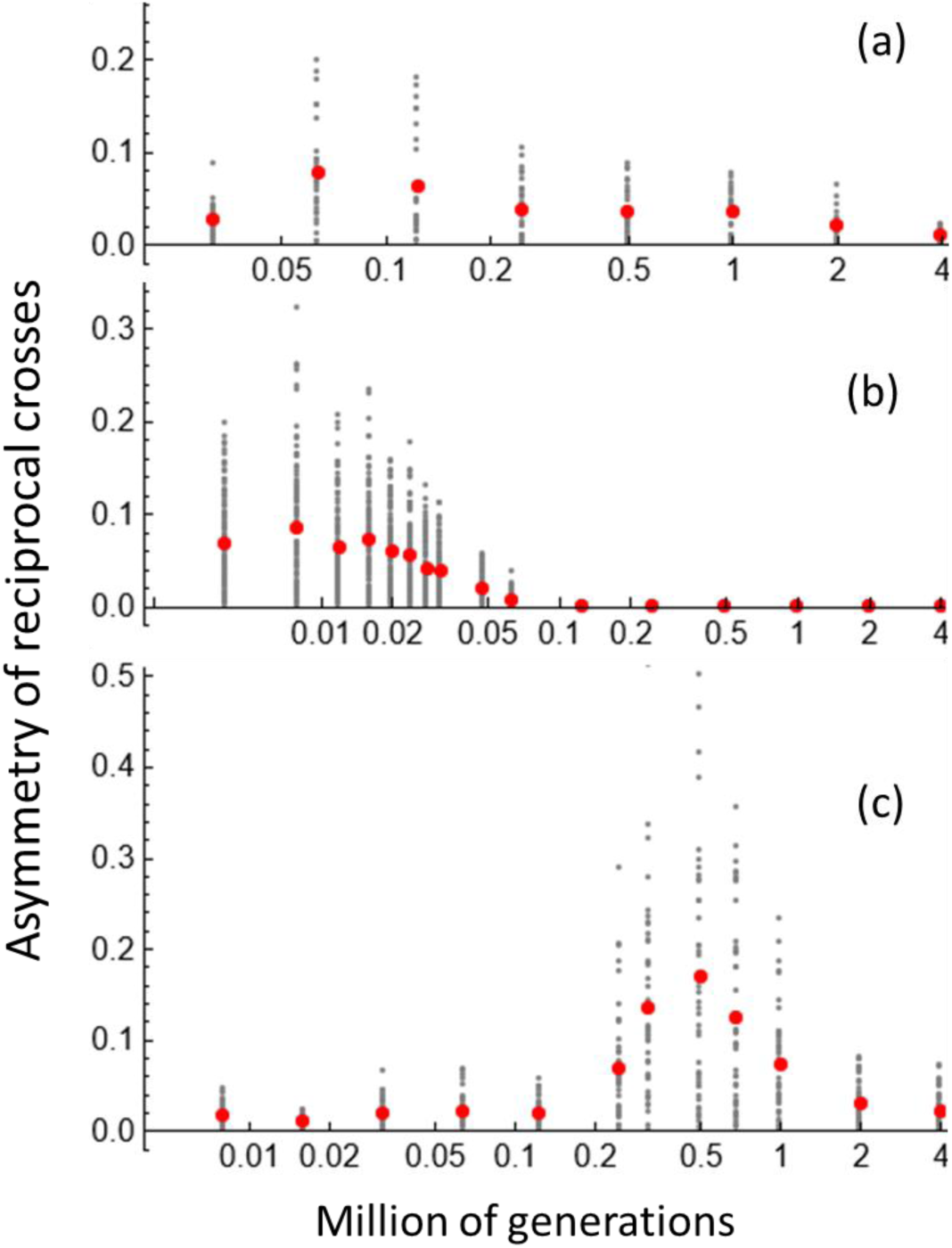
Darwin’s corollary in other scenarios. **(a)** Crosses asymmetries in XO simulations (same simulations as the one illustrated in Fig 2D). The graphs show the absolute value of the difference in the fitness of males produced in reciprocal crosses (y-axes) against time (x-axis, in millions of generations, in log-scale). A larger value means that the male fitness reduction is larger in one direction of the cross. Gray dots are the individual values obtained from all hybrids among independently evolving populations; red dots indicate mean values. **(b)** Same as in (a), but for male-limited simulations with 500 genes (same simulations as the unsorted case illustrated in Fig 2E). (**c**): Same as in (a), but for male-limited simulations with 50 genes. The x-axis does not represent the exact same range of values in the three panels to better emphasize the period during which asymmetries are maximized.

#### 1.7 Numerical examples for the different situations

**Fig S7a.**
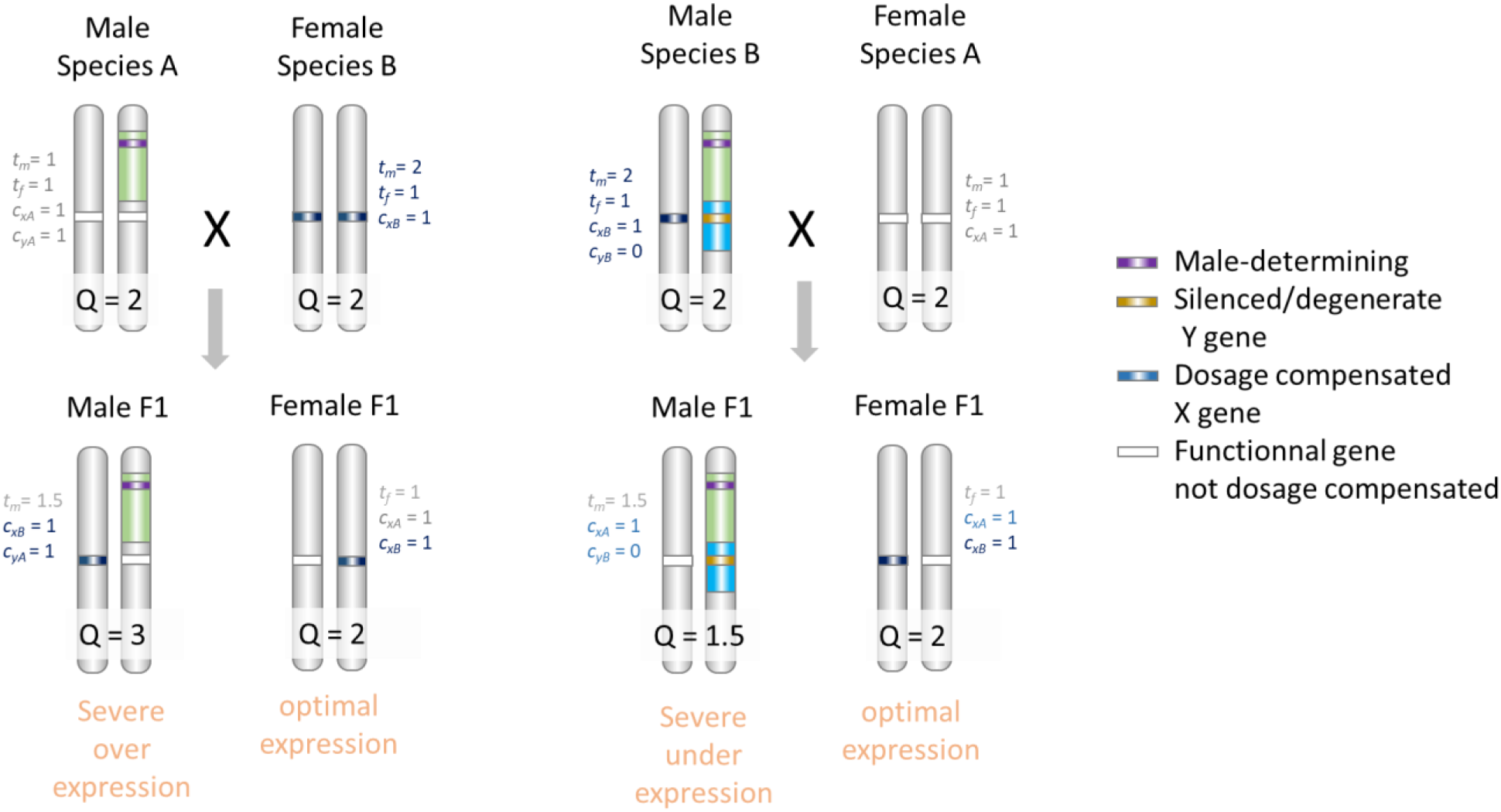
Example of the effect of a gene located in a derived stratum present in only one species, with a drosophila-like DC mechanism. The figure shows species A and B, where the Y has a non-recombining stratum (in green) inherited from the common ancestor, and a newly evolved non-recombining stratum only in species B (in blue), as in Fig S10a. The example focuses on a gene in the derived (blue) stratum. In species A, DC does not evolve (since the gene is located in a recombining portion of the Y), while in species B, DC is achieved for that gene, since it is located in the non-recombining region. In this example, DC in species B is achieved by having male overexpression of the X, like in *Drosophila* (i.e. by evolving a male-specific trans-acting factor increasing X expression in males, *t_m_* = 2 in species B). Male and female expression levels is indicated by the value of *Q*. *Q* = 2 is the optimal expression level (*Q* is computed as (*c*_*X*,*i*_ + *c*_*Y*,*i*_)*t̄*_*m*,*i*_ and (*c*_*X*1,*i*_ + *c*_*X*2,*i*_)*t̄*_*f*,*i*_ in males and females, respectively, see §1.1). F1 males show a departure from optimal expression, while F1 females maintain optimal expression, in both directions of the cross. Note that the reduction in F1 male fitness is potentially asymmetric in the two directions of the cross, resulting from either over or underexpression.

**Fig S7b.**
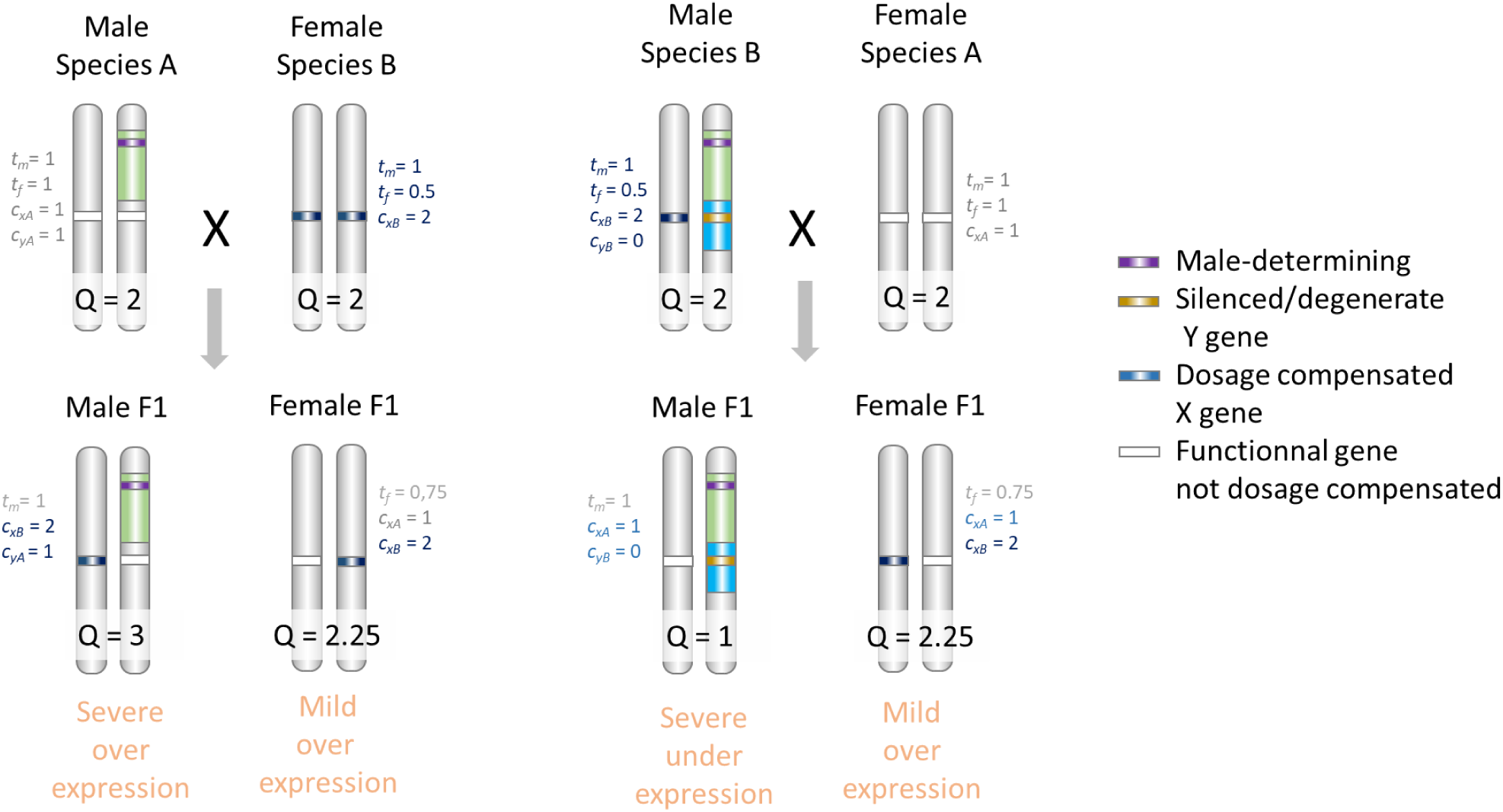
Example of the effect of a gene located in a derived stratum present in only one species, with a mammal-like DC mechanism. The figure shows species A and B, where the Y has a non-recombining stratum (in green) inherited from the common ancestor, and a newly evolved non-recombining stratum only in species B (in blue), as in Fig S10a. The example focuses on a gene in the derived (blue) stratum (like in Fig S7a). In species A, DC does not evolve (since the gene is located in a recombining portion of the Y), while in species B, DC is achieved for that gene, since it is located in the non-recombining region. The example shows a mammal-like pattern of DC for species B: DC in species B is achieved by having overall X overexpression (by evolving stronger X cis-regulatory elements, *c_xB_* = 2 in species B) which is corrected in females to avoid overshooting (by evolving a female-specific trans-acting factor decreasing X expression in females, *t_f_* = 0.5 in species B). Male and female expression levels is indicated by the value of *Q*. *Q* = 2 is the optimal expression level (*Q* is computed as (*c*_*X*,*i*_ + *c*_*Y*,*i*_)*t̄*_*m*,*i*_ and (*c*_*X*1,*i*_ + *c*_*X*2,*i*_)*t̄*_*f*,*i*_ in males and females, respectively, see §1.1). F1 males show a greater departure from optimal expression than F1 females. Note that the reduction in F1 male fitness is potentially asymmetric in the two directions of the cross, resulting from either over or underexpression.

**Fig S8.**
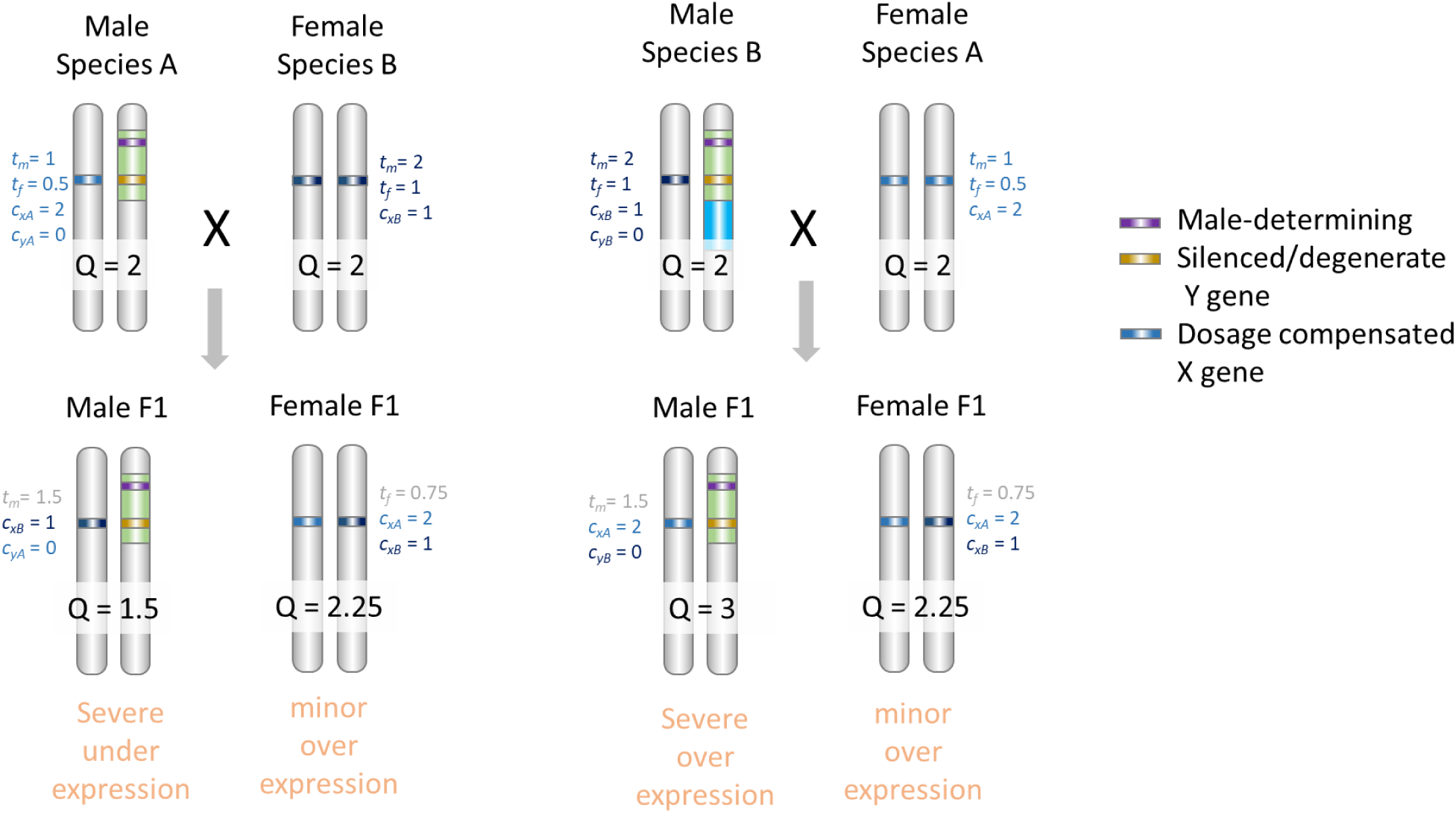
Example of the effect of a gene located in an ancestral stratum, with different DC mechanisms in the diverging species. The figure shows species A and B, where the Y has a non-recombining stratum (in green) inherited from the common ancestor, and a newly evolved non-recombining stratum only in species B (in blue), as in Fig S10a. The example focuses on a gene in the ancestral (green) stratum. In species A, DC is achieved, for the focal gene, by having overall X overexpression (by evolving a stronger X cis-regulatory element, *c_xA_* = 2 in species A) which is corrected in females to avoid overshooting (by evolving a female-specific trans-acting factor decreasing X expression in females, *t_f_* = 0.5 in species A). In species B, DC is achieved, for the same gene, by having male overexpression of the X (i.e. by evolving a male-specific trans-acting factor increasing X expression in males, *t_m_* = 2 in species B). Male and female expression levels is indicated by the value of *Q*. *Q* = 2 is the optimal expression level (*Q* is computed as (*c*_*X*,*i*_ + *c*_*Y*,*i*_)*t̄*_*m*,*i*_ and (*c*_*X*1,*i*_ + *c*_*X*2,*i*_)*t̄*_*f*,*i*_ in males and females, respectively, see §1.1). Male F1 show a greater departure from optimal expression than female F1, in both directions of the cross. Note that the reduction in F1 male fitness is potentially asymmetric in the two directions of the cross, resulting from either over or underexpression.

**Fig S9a.**
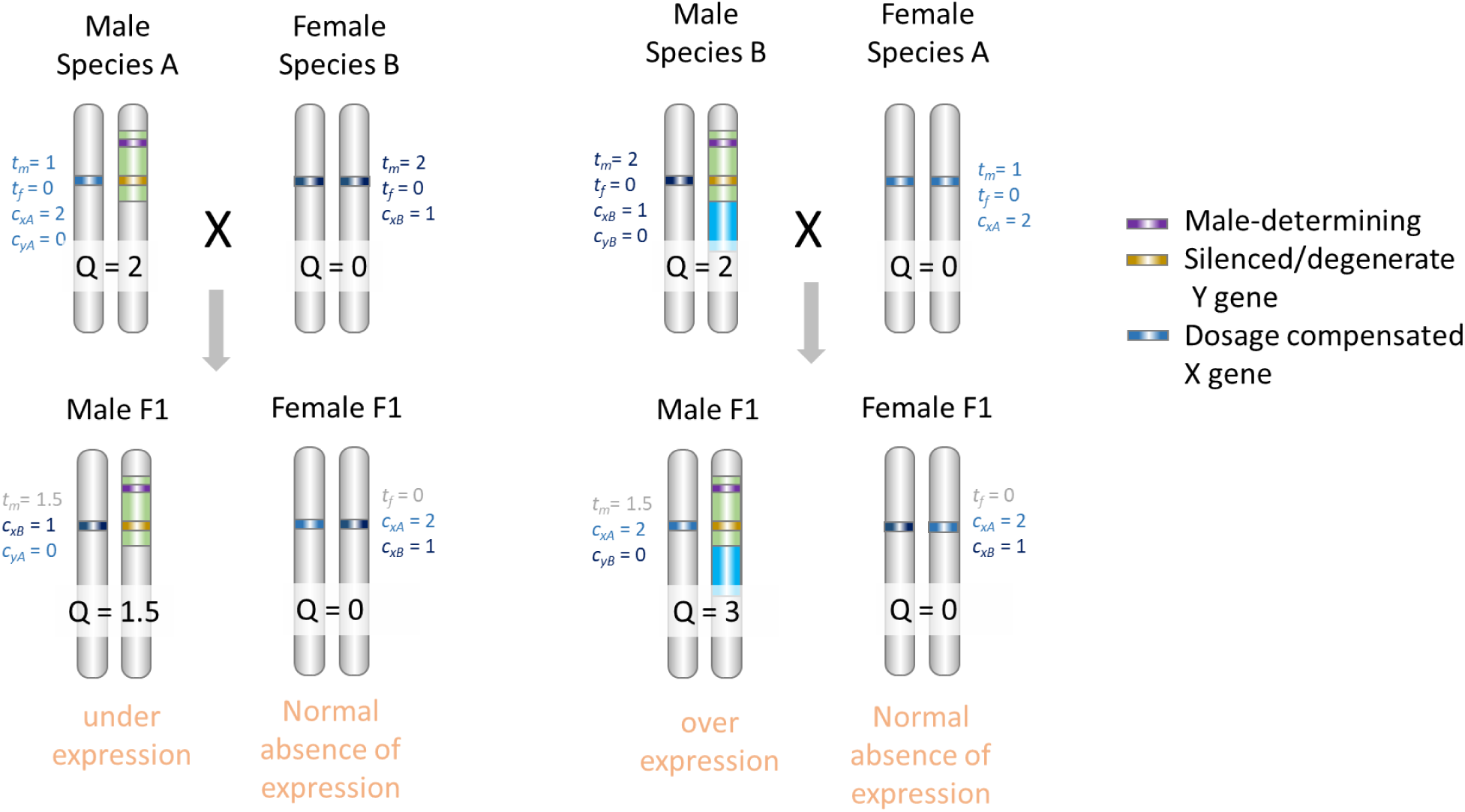
Example of the effect of a male-limited gene located in an ancestral stratum, with different DC mechanisms in the diverging species. The figure shows species A and B, where the Y has a non-recombining stratum (in green) inherited from the common ancestor, and a newly evolved non-recombining stratum only in species B (in blue), as in Fig S10a. The example focuses on a male-limited gene in the ancestral (green) stratum. The gene is not expressed in females. In species A, DC is achieved, for the focal gene, by having overall X overexpression (by evolving stronger X cis-regulatory elements, *c_xA_* = 2 in species A) which is corrected in females to avoid overshooting (by evolving a female-specific trans-acting factor decreasing X expression in females, *t_f_* = 0.5 in species A). In species B, DC is achieved, for the same gene, by having male overexpression of the X (i.e. by evolving a male-specific trans-acting factor increasing X expression in males, *t_m_* = 2 in species B). Male and female expression levels is indicated by the value of *Q*. *Q* = 2 is the optimal expression level (*Q* is computed as (*c*_*X*,*i*_ + *c*_*Y*,*i*_)*t̄*_*m*,*i*_ and (*c*_*X*1,*i*_ + *c*_*X*2,*i*_)*t̄*_*f*,*i*_ in males and females, respectively, see §1.1). Male F1 show a departure from optimal expression in both directions of the cross. Note that the reduction in F1 male fitness is potentially asymmetric in the two directions of the cross, resulting from either over or underexpression. Note also that this situation is identical to Fig S7, ignoring females.

**Fig S9b.**
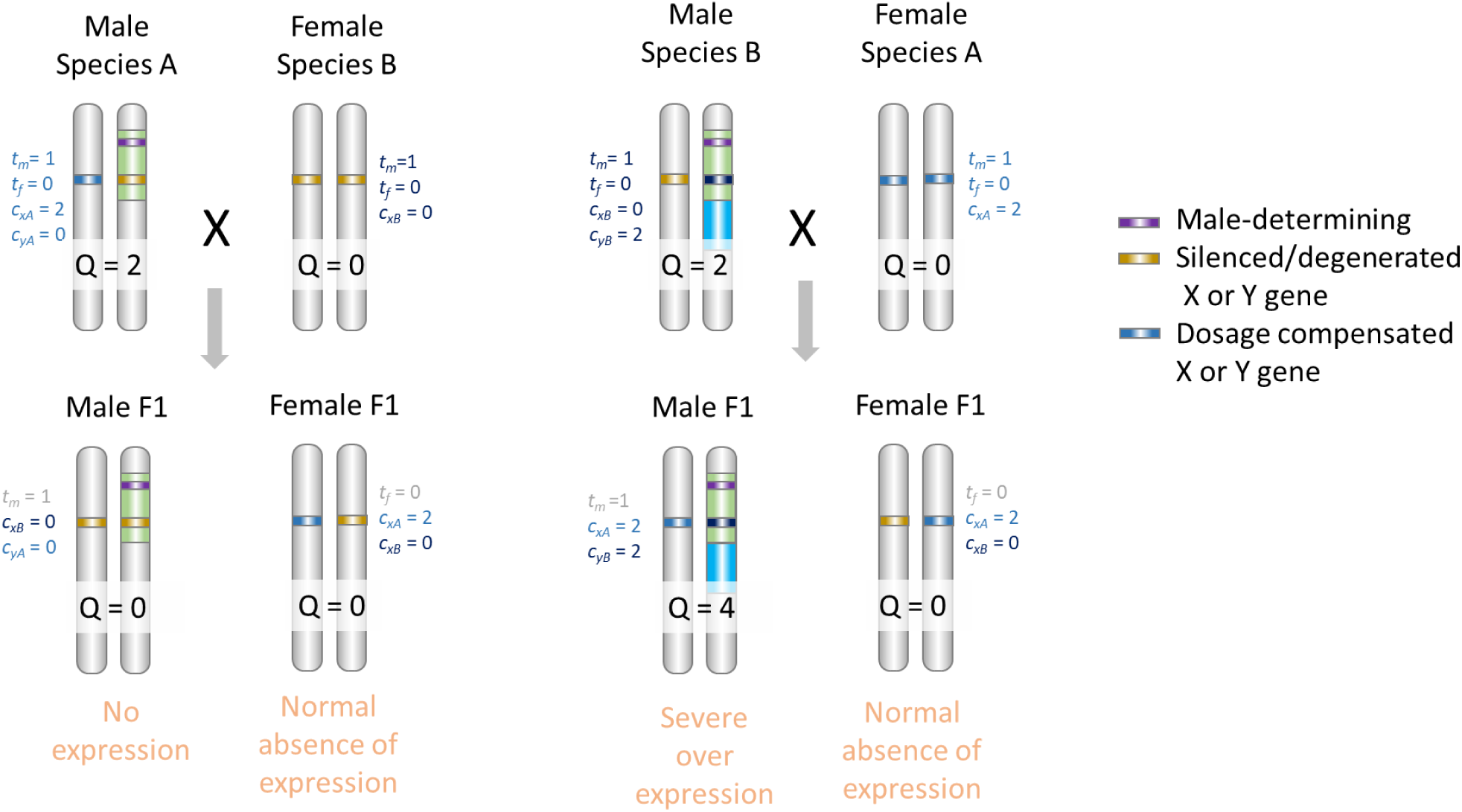
Example of the effect of a male-limited gene located in an ancestral stratum, with alternate X or Y degeneration in the diverging species. The figure shows species A and B, where the Y has a non-recombining stratum (in green) inherited from the common ancestor, and a newly evolved non-recombining stratum only in species B (in blue), as in Fig S10a. The example focuses on a male-limited gene in the ancestral (green) stratum which is silenced on the Y in species A (*c_yA_* = 0), but silenced on the X in species B (*c_xB_* = 0). Indeed, male-limited genes can degenerate both ways, which differs from non-sex-limited cases shown in Fig S7 and S8 where degeneration can only occur on the Y. The gene is not expressed in females. In both species, DC is achieved by having male overexpression of either the X or Y, through the evolution of stronger cis-regulators (*c_xA_* = *c_yB_* = 2). Male F1 show a strong departure from optimal expression in both directions of the cross, notably in one direction where the gene is completely silenced. Note that the reduction in F1 male fitness is potentially asymmetric in the two directions of the cross, resulting from either over or underexpression.

**Fig S9c.**
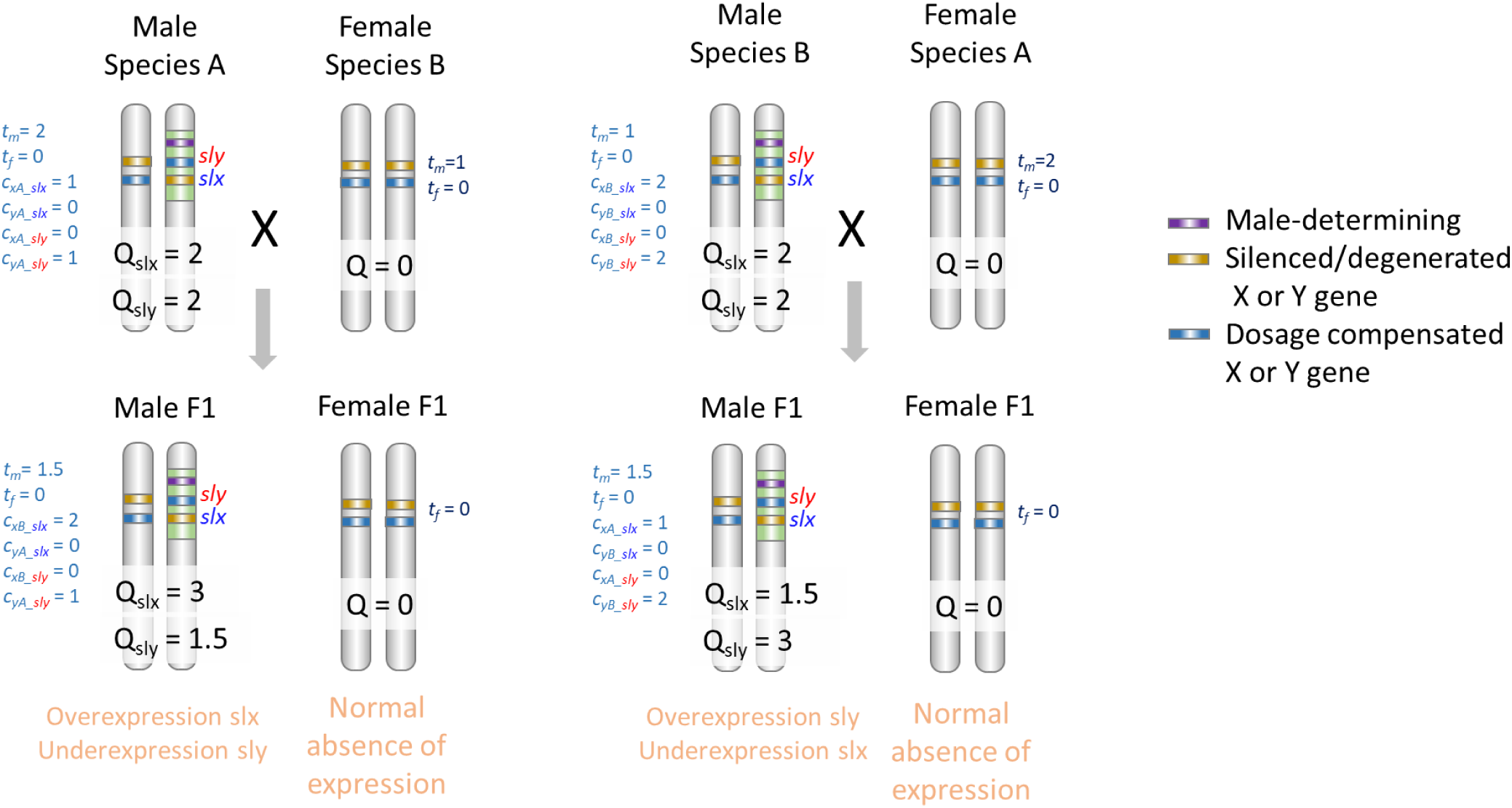
Example of sly/slx; discussed in section 2.4. The figure shows species A and B, where the Y has a non-recombining stratum (in green) inherited from the common ancestor. Gene *slx* is only present on the X (the Y copy is degenerate), while gene *sly* is only present on the Y (the X copy is degenerate). Both *slx* and *sly* are male limited genes, only expressed in the male germline, with optimal expression level *Q*_opt_ = 2. This is aimed at representing a simplified situation close to that of *slx* and *sly* genes in mice (*102*). These genes are present in multiple copies on the X or Y, but with a different number of copies in different species. This is equivalent in our model to having stronger of weaker cis-regulators (having multiple copies of a gene on a non-recombining part of the sex chromosomes is equivalent, here, in terms of expression of having a single copy with a stronger cis-regulator). In species A, *slx* and *sly* have a low copy number, and this is represented by a low value for their cis-regulator (c_XA_slx_ = c_YA_sly_ = 1). Assuming they are interacting with the same trans-acting factor (only expressed in male), the strength of this trans acting factor is such that it ensures optimal expression of both genes (*t*_m_ = 2). In species B, *slx* and *sly* have a large copy number, and this is represented by a larger value for their cis-regulators (c_XB_slx_ = c_YB_sly_ = 2). Assuming for simplicity they are interacting with the same trans-acting factor (only expressed in male), the strength of this trans acting factor is such that it ensures optimal expression of both genes (*t*_m_ = 1). As the figure illustrates, depending on the direction of the cross, male F1 will either exhibit overerexpression of *slx* and underexpression of *sly*, or the reverse. The fitness consequences of these departures will depend on the exact phenotypic effect of each gene. The fitness of F1 males will also be reduced if fitness is decreased by an unbalance of *slx* and *sly* expression.

### 2 Computing hybrid fitness

The decrease in fitness resulting from the departure from optimal dosage *W*^*Q*^ depends on the intensity of stabilizing selection on the expression level, *I* (Eq. 4). Eq. 4 ensures that complete loss of expression (at *Q* = 0) has the same fitness effect as the fitness effect of completely disrupting the coding sequence (*s_max_*, see Methods). For the purpose of this analysis, we assume that *Q*_*opt*_= 2 without loss of generality. We also define *θ* = *I s*_*max*_, which represents the overall intensity of stabilizing selection on dosage. We drop below the indices *i* referring to a specific gene, to simplify the notation.

A variety of of dosage compensation mechanisms have been described across a range of organisms, which may be represented by specific instance of our general model. Indeed, once the Y is fully silenced (*c*_*Y*_ = 0), and assuming that the population remains at *Q*_*opt*_ in both males and females (which is a good approximation unless stabilizing selection is particularly weak), we have

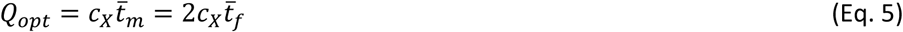

We therefore have *t̄*_*m*_ = 2*t̄*_*f*_, which defines dosage compensation. Hence the manner in which DC operates (i.e. the triplet *t̄*_*m*_, *t̄*_*f*_, *c*_*X*_) can be delineated by a single parameter, given that there is only one degree of freedom in the way DC occurs. For the purpose of this description, we may, for instance choose to use *t̄*_*m*_. In comparison to the initial system in which *t̄*_*m*_ = *t̄*_*f*_ = *c*_*X*_ = *c*_*Y*_ = 1, a final compensation characterized by *t̄*_*m*_ =1 (i.e. *t̄*_*m*_ = 1, *t̄*_*f*_ = 0.5, *c*_*X*_ = 2, *c*_*Y*_ = 0 ) would correspond to the *Caenorhabditis elegans* case, where the X is inherently expressed twice as much (*c*_*X*_ = 2) to achieve optimal expression in males, while an hermaphrodite-limited *trans-*regulator halves expression (*t̄*_*f*_ = 0.5) to recover optimal expression in hermaphrodites. This is also analogous to the mammal case where a female-limited *trans-*regulator halves expression by randomly silencing one X (instead of halving the expression of each X like in *C. elegans*). The case *t̄*_*m*_ = 2 (i.e. *t̄*_*m*_ = 2, *t̄*_*f*_ = 1, *c*_*X*_ = 1, *c*_*Y*_ = 0 ) would correspond to the *Drosophila* case, where a male-limited *trans-*acting factor doubles X expression to achieve optimal expression in males (with no change in females, *t̄*_*f*_ = 1, *c*_*X*_ = 1).

#### 2.1 Contribution of autosomes to the fitness reduction of F1 hybrids

On autosomes, cis and trans regulators can coevolve provided that the overall expression level stays close to its optimum value. This condition implies that, for each gene, *t*_*m*_ and *t*_*f*_ stay both close to 1/*c.* Hence, autosomes can contribute to a reduction in hybrid fitness, but it will be symmetrical for males and females. Noting Δ the difference in *t* values between the two species for a given gene, and assuming that this difference is weak, we find, from Eq. 4, that the reduction in fitness of F1 hybrids caused by misregulation of this gene is, to leading order in Δ:

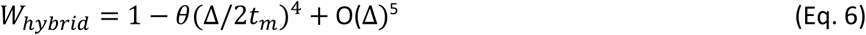

This effect is modest (note the quartic exponent), but can lead to substantial fitness loss if cumulated over many genes. The requirement of cis and trans regulators to match is a particular case of a signal-receiver or key-and-lock type of situations. Regulators can relatively “freely” evolve as long as they match (*98*), potentially creating postzygotic isolation after divergence throughout the genome.

#### 2.2 Contribution of ancestral and derived Y strata to the fitness reduction of F1 hybrids

We can compute the male/female fitness ratio in hybrids, accounting for two types of loci (Fig S10a). We consider first the Y-linked loci that are degenerate and silenced in both species (assume there are *k*_1_ such loci). These loci are for instance located in a non-recombining region that is common to both species (green stratum 1 in Fig S10a, where the sex-determining locus, SDL, is located). Second, we consider loci that are degenerate and silenced in only one species (assume there are *k*_2_ such loci). These loci are for instance located in a non-recombining region that only evolved in one species (blue stratum 2 of species A in Fig S10a). We compute the contribution to Haldane’s rule of these two types of loci in turn.

**Fig S10a.**
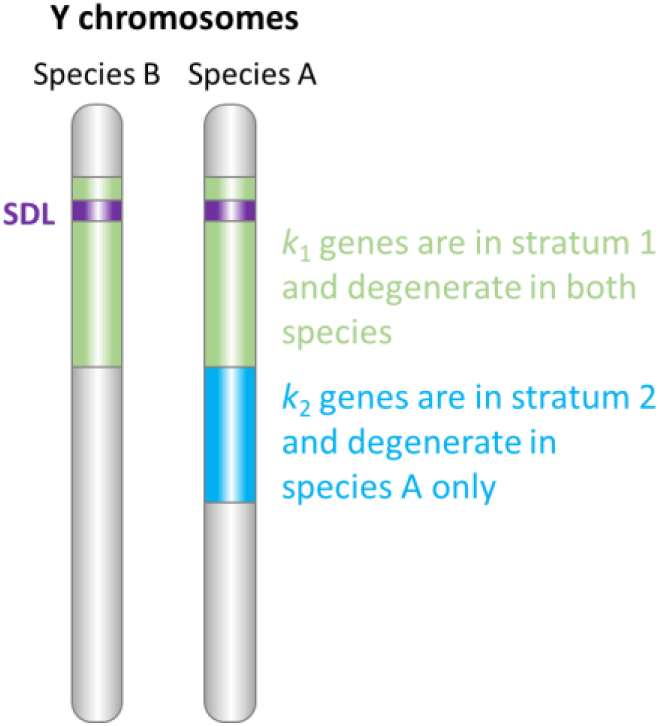

##### 2.2.1 Contribution of an ancestral stratum (stratum 1)

For loci that are degenerate in both species, the contribution of a given gene to the male and female fitness can be approximated assuming that the gene is fully silenced on the Y and dosage-compensated in both species, that is, without loss of generality *c*_*Y*_ = 0, *c*_*X*_ = 2/*t*_*m*_, *t*_*f*_ = *t*_*m*_/2. Noting Δ the difference in *t*_*m*_ values between the two species for a given gene, and assuming that this difference is weak, we find, from Eq. 4, that the contribution to Haldane’s rule of that gene is, to leading order in Δ:

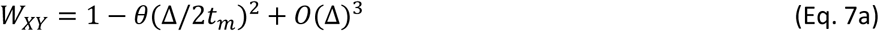

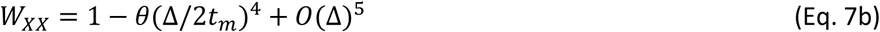

For a given divergence, the fitness reduction in males is much larger than in females (which equals the fitness reduction expected for an autosomal locus). This is due to an averaging mismatch: in males, a single functional cis-regulator is inherited on the maternal X (the cis-regulator on the Y is silenced), but trans-regulators are inherited from both parents. The male to female fitness ratio *ρ* is:

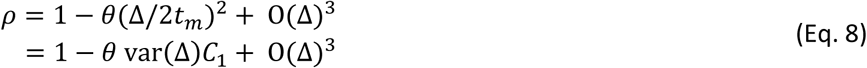

where var(Δ) is a measure of the divergence in the DC mechanism between the hybridizing species, equal to E(Δ^2^), and *C*_1_ a positive constant equal to the inverse of 4*t*^2^ .

##### 2.2.2 Contribution of a derived stratum (stratum 2)

The male/female ratio contributed by the loci in stratum 2 can be approximated assuming that in species B the Y genes are not degenerate and remain fully expressed (i.e. *t̄*_*m*_ = *t̄*_*f*_ = *c*_*X*_ = *c*_*Y*_ = 1), while in species A the genes are degenerate, silenced, and dosage compensated (with a given *t*_*m*_ value, i.e. without loss of generality *c*_*Y*_ = 0, *c*_*X*_ = 2/*t*_*m*_, *t*_*f*_ = *t*_*m*_/2). The reciprocal crosses are asymmetrical for males, depending on which Y they inherit. In cross 1 (male A x female B), the Y in hybrid male offspring is carrying degenerated genes, but in cross 2 (male B x female A), it does not.

Assuming weak stabilizing selection, we find, for the effect of one gene:

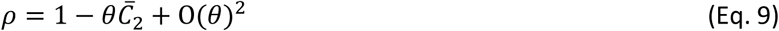

Where *C*_2_ is a constant equal to *C*_2*a*_ in cross 1, and to *C*_2*b*_ in cross 2 and to *C̄*_2_ = (*C*_2*a*_ + *C*_2*b*_)/2 on average over the two types of crosses, with:

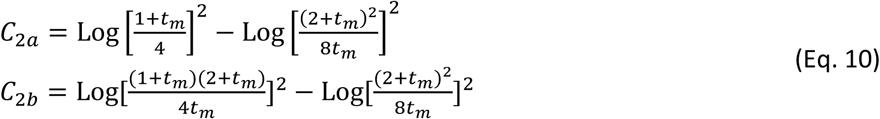

*C*_2*a*_ and *C*_2*b*_ are positive (except in a very small region around *t*_*m*_=3 where *C*_2*a*_is very close to zero, Fig S10b). *C*_2*a*_ could also be negative for very low values of *t*_*m*_ (*t*_*m*_ < 0.17), but these situations are probably not relevant as they imply varying cis regulator strength by an order of magnitude, and are far outside values consistent with extent DC systems (ie. *t*_*m*_ between 1 and 2). Fig S10b shows the value of these constants in the plausible range 0.5 < *t*_*m*_< 4.

**Fig S10b.**
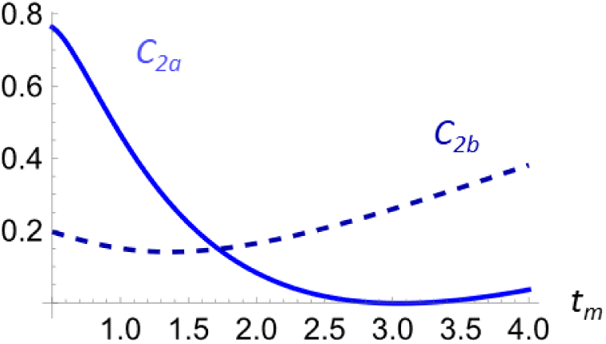

##### 2.2.3 The overall effect of ancestral and derived strata

Hence with weak stabilizing selection and at small divergence, the male/female fitness ratio among F1 offspring, overall, is to leading order close to

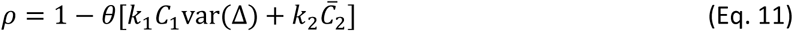

With *k*_1_ genes on the shared ancestral stratum and *k*_2_ genes in the recently evolved stratum in one of the two species, where *C*_1_ and *C̄*_2_are positive constants that depend on the DC mechanism, and var(Δ) is a measure of the divergence in the DC mechanism between the hybridizing species. Hence, HR due to the misregulation of dosage compensated genes can be caused by the portion of the Y that is degenerate and compensated in both species (provided they exhibit some divergence in DC), and by the portion of the Y that is degenerate and compensated in only one of them. Importantly, the fitness effect in hybrids directly scales with the size (i.e. the number of genes) of the non-recombining strata (shared or recently evolved), everything else being equal. However, this is not strongly discriminating with other theories. Such scaling is also expected, for instance, for the dominance or the meiotic drive theory.

In a drosophila-like situation where dosage compensation is achieved by overexpressing the X in males (*t*_*m*_ around 2), we have:

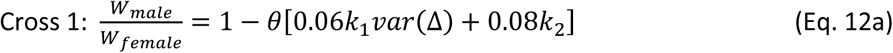

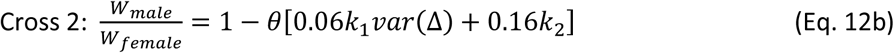

while in a *C. elegans* or mammal case (*t*_*m*_ around 1):

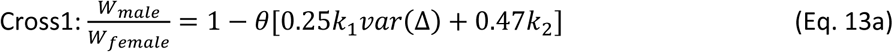

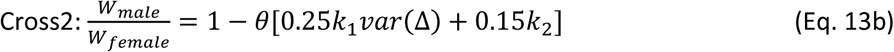

In all cases, Haldane’s Rule occurs and combines a regulatory effect of the genes that are degenerate in one species and not the other, and a regulatory effect of the genes that are degenerate in both species (but not compensated exactly in the same way in the two species). The two effects combine and contribute to HR.

Loci that are degenerate in one species but not in the other generate an expression mismatch in hybrids, even if trans-acting factors are codominant. In the range of known DC mechanisms, i.e. with *t*_*m*_ values between 1 and 2, hybrid females will tend to show slight overexpression, males from cross 1 will show relatively severe underexpression, and males from cross 2 relatively severe overexpression (Fig S7-S9).

For loci that are degenerate in both species, the effect comes from the fact that genes may exhibit a quantitative difference in the way they are dosage compensated (since achieving optimal DC can be obtained by a different combination of cis and trans effects, i.e. different *t*_*m*_ values). This difference will be however more buffered in females (where cis effects are averaged over the two X) than in males (where cis effects are not averaged and only expressed from a single X, Fig S8). In both males and females, (autosomal) trans-effects are equally averaged. In species pairs sharing an identical and chromosomal-level DC mechanism, this effect is expected to be relatively minor, since genes should not exhibit quantitative differences in the DC mechanism. However, in many species, many genes escape this global DC mechanism and exhibit gene-level expression control (*103–109*). These genes, if sufficiently numerous on the X could therefore also contribute to HR. Current evidence indicates that chromosome-wide DC is the exception, rather than the norm (*59–61*).

Finally, in systems where DC occurs around a Drosophila-like situation (*t*_*m*_ around 2), hybrid males inheriting the most degenerated Y (from cross 1) suffer more than males inheriting the less degenerated Y (from cross 2). The reverse occurs in systems where DC is close to a C. elegans/mammal type (*t*_*m*_ around 1). However, the difference will manifest only in species pairs showing Y with different degrees of degeneration (i.e. only if *k*_2_>>1 and larger than *k*_1_).

#### 2.3 Case of unbalanced females

In several Drosophila experiments, the fitness of F1 females carrying 2 attached X from the same species (XXY females) was investigated to test the dominance theory of Haldane’s rule. In these cases, it was expected that homozygous females (carrying two X from the same parental species) would show a large fitness reduction, like hemizygous males (but unlike standard heterozygous XX females). These crosses have revealed that when HR was about F1 fertility, unbalanced females remained fertile (unlike F1 males), while when HR was about F1 viability, unbalanced females were inviable (like F1 males). These results generated considerable discussion (*1*, *13–16*, *99*, *110*, *111*).

Using the same approach as above, and assuming perfect dosage compensation in the parental species, we can first compute the expected fitnesses of the different F1s for genes located on stratum 1 (shared between the two hybridizing species), assuming a small divergence in their regulatory traits (small Δ) :

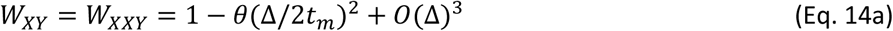

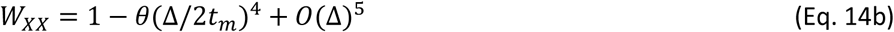

For genes in that stratum, we find that unbalanced females present the same fitness reduction as males, much larger than the one seen in regular F1 females. For genes located on stratum 2, degenerate and compensated only in species A, and noting X^d^ the X from species A, we find that the fitness reduction is proportional to *θK*, where *K* is a constant that depends on how X^d^ is dosage compensated (i.e. it depends on the value of *t*_m_). The different values of *K* are straightforward to compute and are illustrated in Fig. S10c for the different F1 in the reciprocal crosses. Unbalanced females have a lower fitness than standard females except near *t_m_* =2 where the difference is small. Overall, combining effects in stratum 1 and 2, we can therefore conclude that this model predicts that unbalanced F1 females show a greater fitness reduction compared to standard females. The comparison to males depends on whether genes in stratum 1 or 2 contribute. For genes on stratum 1, the fitness reduction is the same as in males. For genes on stratum 2, it depends on the direction of the cross and *t_m_* values (see Fig. S10c). Overall, these results indicate that unbalanced F1 females should often present a large reduction in fitness, much more similar to that of F1 males than that of F1 females. This result holds for viability. For sterility, the fitness of males and females (balanced or unbalanced) will differ, if genes involved in sterility are expressed in a sex-specific manner (see next section for the effect of such genes). As shown below, male-limited genes will cause a fitness reduction in F1 males, but female-limited genes will not cause a fitness reduction in F1 females. This would therefore explain the contradictory results observed for the *Drosophila* experiments mentioned above where unbalanced females are fertile when HR involves sterility but are inviable when HR involves viability (*15*, *110*).

**Fig S10c.**
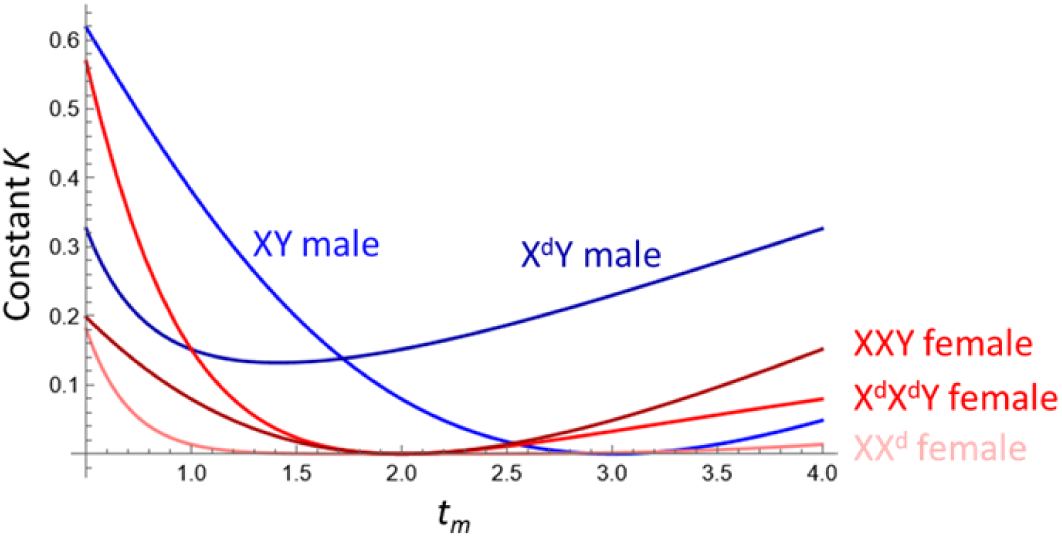

#### 2.4 Case of sex-limited genes

We can consider the case of fertility genes. We consider that these genes are expressed in only one sex (they are e.g. expressed in the germline), contrary to viability genes, which in the vast majority of cases will be expressed in both sexes). We will take male as the heterogametic sex as in XX/XY species, but the same argument can be made by switching sexes for ZZ/ZW species.

Female-limited genes can obviously be lost from the Y, as they are not expressed in males. Apart from this, their regulators can diverge between species, like in the case of autosomes, generating a consistent but small decrease in fitness in F1 (as in Eq. 6). Given the lower effective population size on the X (compared to autosomes), and the weaker selection pressure overall (everything else being equal, sex-limited genes are only selected in one sex), we might nevertheless expect that the regulatory divergence could somehow be faster on the X compared to autosomes (as seen in the results, compare the female fitness drop illustrated on Fig 2E to that on Fig 2A).

Male limited genes can be lost from the Y (like genes expressed in both sexes), but they can also be lost from the X (see (*112*) and empirical example in (*113*, *114*)). Indeed, there is no requirement that they remain functional in XX females (contrary to genes expressed in both sexes that can only evolve Y silencing (*36*)). Diploid expression is unstable after recombination arrest, but silencing can occur both ways. Simulation results confirm this finding. However, despite this apparent symmetry, X and Y silencing is not occurring at an equal rate. Male-limited genes are more likely to be silenced on the Y than on the X (and indeed such cases are documented (*115–117*). Note that it does not contradict the observation that most genes remaining functional on the Y are male limited. Only those genes can be maintained on the Y (while degenerating on the X). The bias is for instance close to 6:4 in a non-recombining stratum with 50 male-limited genes, but it is 9:1 in a stratum with 500 male-limited genes (in both cases the simulation had a population size of 10,000 individuals, and the X or Y silencing occurred relatively quickly). Presumably, the bias increases with the relative ease at which deleterious mutations initially accumulate on the Y versus X once recombination has stopped (due to selective interference), preferentially pushing the regulatory feedback loop in that direction. In all cases, because one copy is silenced, these male-limited genes will evolve dosage compensation to stay expressed at the right level in males. Their cis-regulators on the X or Y will coevolve with the trans-regulator expressed in males. Note that having multiple copies of a gene on a non-recombining part of the Y is equivalent in terms of expression of having a single copy with a stronger cis-regulator. Hence the observation of frequent duplication of Y male-limited genes may correspond in part to the evolution of dosage compensation for those genes. Note too that the simple example given above can extend to more intricate situations, for instance when male limited genes reciprocally lost on the X and Y interact. A case in point is the slx/sly system in mice (*102*). Fig S9c illustrates how such situations could relate to our theory.

In an F1 hybrid, when the focal gene has been silenced on the same chromosome in both species, it will therefore cause a reduction in fitness that is similar to the case of a gene expressed in both sexes (i.e. Eq. 11 applies). If *p* is the chance that a gene is silenced on the Y, and 1 − *p* on the X, this situation has *p*^2^ + (1 − *p*)^2^ chances to occur. If the focal gene is silenced on the Y in one species, but on the X in the other, male F1 will however suffer from a larger fitness reduction. In one direction of the cross, the F1 male will receive two silenced copies, and its fitness will therefore be reduced by a factor 1 − *s*_*max*_. This has *p*(1 − *p*) chances to occur. For an essential gene involved in male fertility, a single case like this would be sufficient to cause complete male sterility. In the other direction of the cross, the F1 male will receive two non-silenced copies, which will correspond to severe overexpression. Its fitness will be reduced by a factor (1 − *θ* log(2)^2^), which is close to (1 − *θ*/2). This has also *p*(1 − *p*) chances to occur. Simulations starting with an XY non recombing pair with 500 fully expressed male-limited genes leads to a dramatic reduction of F1 male fitness, due to this sorting effect (Fig 2E, unsorted). Some genes become silenced on the X and some on the Y, but they are not the same in the two diverging species, resulting in many mismatches (a similar sorting effect can occur in asexual genomes (*118*)). Note that genes are not equally likely to degenerate on the Y and on the X. The former is much more likely (Fig S11), consistent with the idea that the lower effective population size of the Y and the absence of recombination and selective interference on the Y bias silencing and degeneration of the Y copy in this, otherwise, symmetrical situation.

A simulation starting with 500 male-limited genes that are already sorted leads to an F1 male fitness reduction closer to that observed for the XO simulations with genes expressed in both cases. The effect is stronger, however (compare Fig. 2D and 2E), as male-limited genes silenced on the X (and maintained on the Y) will exhibit faster regulatory divergence, due to their lower effective population size (3 times lower than that of genes silenced on the Y and maintained on the X). The initial sorting in these simulations was done following even/odd positions: on even positions 2, 4, …, 498, 500 the gene was initially degenerate (*s* = *s_max_*), silenced on the Y (*c*_Y_ = 0) and dosage compensated (*c*_X_ = 1, *t*_m_ = 2, t*_f_* = 1) ; on odd positions 1, 3,…,499, the gene was initially degenerate (*s* = *s*_max_), silenced on the X (*c*_X_ = 0) and dosage compensated (*c*_Y_ = 1, *t_m_* = 2, *t_f_* = 1).

Overall, we thus predict a stronger effect on HR of male-limited genes than of genes expressed in both sexes. The argument was made for males but would apply to females in ZZ/ZW species. This finding indicates that our regulatory theory could account for the importance of cases of sterility in HR, without having to suppose a ‘faster male’ theory or rely on the systematic presence of divergent meiotic drive systems in sister species. It also better explains why HR is not weaker in Lepidoptera where males are homogametic [a difficulty of the faster male theory noted by Presgraves (*100*)].

**Fig S11.**
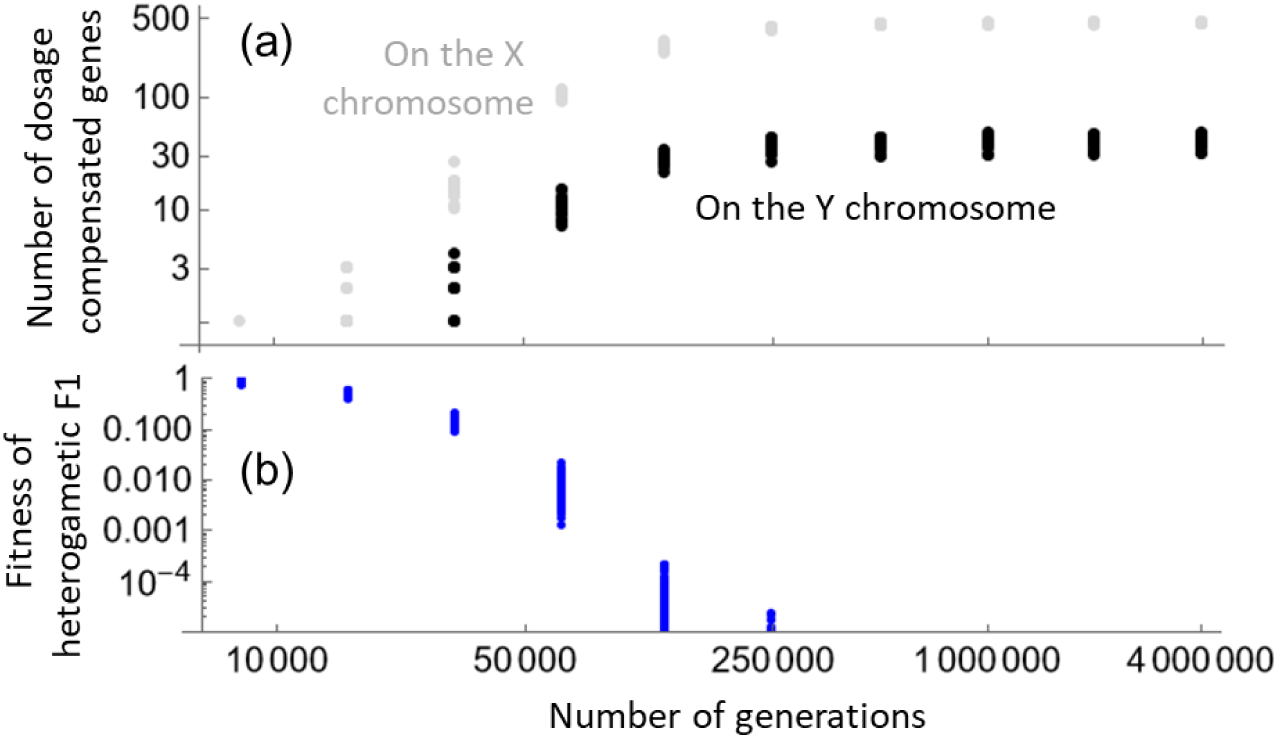
(a) Relative number of male-limited dosage compensated genes evolving on the X and Y chromosomes. In the “unsorted” simulations shown on Fig 2E, among the 500 fully functional male-limited genes initially present on the X and Y, some degenerate on the Y (and stay on the X and become dosage-compensated, gray dots), and some degenerate on the X (and stay on the Y and become dosage compensated, black dots). The figure shows how many genes (y-axis in log-scale) end up in each category through time (x-axis in generations, in log-scale). Despite the symmetry of the regulatory instability, many more genes degenerate on the Y, probably due to selective interference on the non-recombining Y and lower effective population size of the Y. (b) Corresponding fitness of the heterogametic F1 (y-axis, log scale) through time (x-axis in generations, in log-scale). The fitness decrease is fast due to the independent degeneration of genes on the X or Y in different species.

#### 2.5 X-autosomes versus Y-autosomes hybrid incompatibilities

The issue of a possible role of Y-autosome or X-Y interaction for Haldane’s rule has been proposed by Muller (*20*) and often discussed. Several experiments showed that these interactions occur sometimes and contribute to HR on sterility (*119–122*). However, they are far from being the norm (*123*), and are not an explanation for Haldane’s rule on viability (since Y chromosomes have usually very limited effect on viability) or Haldane’s rule in XO species lacking a Y. Most work on HR has therefore mainly focused on X-autosomes interactions.

Most of our results rely on such X-autosomes interactions. Specifically, they concern the misregulation of genes on the X, in crosses between species where at least one evolved silencing and degeneration of those gene copies on the Y. However, it is useful to point out that in some F1 hybrids, the misregulation may involve the non-silenced and non-degenerate Y copy of the gene inherited from the species in which this gene is still on a recombining region of the Y (see e.g. Fig 7a). In such a case, overexpression in F1 males results from the presence of two fully functional copies of the gene (on both the X and Y), in presence of partial dosage compensation (inherited from the species where the Y copy is silenced). In this case, the X and Y gene copies play symmetrical roles, and arguably, this is not best described as a simple “X-autosome” interaction. The fitness reduction involves the autosomal trans-acting factors, and both X and Y copies of the gene. However, the misregulation emerges because in one species, the Y copy of the gene was silenced and the remaining X copy evolved DC. In this sense only can it be presented as a problem of X-linked dosage compensated genes.

However, as simulations involving genes with sex-limited expression show, sometimes genes can also degenerate on the X (or Z in ZW species). Example of genes lost on the X and with a local DC have been reported (*124*). In this case, the dosage compensated (functional) genes may be located on the Y (or W), and their misregulation could also contribute to explain HR on sterility (see previous section). In the simulations illustrated on Fig 2E, unsorted case, 10% of genes stay functional on the Y and degenerate on the X (in a ZW system, similarly, a handful of female-limited genes would stay on the W and degenerate on the Z). These genes would contribute to HR through Y-autosomes interactions. The reason these genes contribute to HR is the same than for other sex-linked genes in our model: the heterogametic sex will average autosomal trans-regulatory effects but not cis Y or W regulatory effects (since a single cis functional sequence is inherited in the heterogametic sex). Hence, our model may explain why Y / W linked genes could contribute to HR on sterility, and why such cases are likely to be rare.

The argument about these Y effects could be somehow generalized. Our simulations concern genes present on the ancestral autosomal pair and degenerating either on the X or on the Y. Yet, the regulatory averaging imbalance between the homo and heterogametic sex in F1 hybrids could also, in principle, apply to genes evolving *de novo* on the Y or the X. As long as the regulation of expression is determined by the interaction between a sex-linked (X or Y) cis-acting factor and an autosomal trans-acting factor, then misregulation is likely to occur more strongly in the heterogametic F1, due to the difference in averaging between cis and trans effects in hetero and homogametic sex. A particular and noteworthy case could involve sequences evolving on the Y (and absent on the X) and acting as ‘chromatin sinks’ (as reported in several studies (*125*)) for an autosomal transcription factor or any autosomal factor that affects chromatin state. Such sequences could have evolved due to an accumulation of transposable elements on the Y, or to regulate specific male-limited genes as often reported (*126–129*). The strength of a such a chromatin sink could be modeled in the same way as a cis-regulatory strength. Consider, for instance, a species without this chromatin sink on the Y. In that species, the autosomal factor is free to interact with its normal target elsewhere in the genome. Consider now another species in which a sink sequence has evolved on the Y. Because this sink sequence “absorbs” some of the autosomal factor, it is less available for its normal function, which presumably would be detrimental. Hence, we would expect that, in this second species, this autosomal factor has evolved to become overexpressed to compensate for this sink effect (and reach the proper amount of factor not captured by the sink sequence). F1 males, however, will inherit either the Y with or without the sink sequence (depending on the cross direction), but always an “averaged” production of the autosomal factor. It will always result in over or underexpression of that factor in F1 males. In F1 females, however, no such effect will occur (since in this example the sink effect is specific to males having a Y with the sink sequence). Hence, it is theoretically and empirically possible that the effects we describe on genes extends to any cis-regulatory element on the X or Y, not just coding genes and their cis-regulatory sequences that were inherited from the ancestral autosomal pair. In particular, all Y/W specific sequences interacting with autosomal factors could possibly play a role in HR in our theory, including sequences acting as chromatin sinks.

### 3 Comparison to other theories

#### 3.1 The many theories for Haldane’s rule and the large X effect

Many theories for HR and LX have been proposed over the years. Not all have been exposed in the main text and in Table 1. In particular many ideas have been already convincingly discarded in the literature. Regarding HR, (*6*) lists and discuss them. For instance, the “sexual transformation”, the “chromosomal rearrangement”, or the “Y incompatibilities” were historically proposed, but can all be convincingly ruled out. Regarding LX, (*11*) also list proposed theories. Obvious possibilities, such as a greatest concentration of sterility genes on the X or Z, have been discarded. Similarly the “Faster X” theory has been theoretically worked out and empirically investigated, but seems quantitatively unlikely to explain the observations (see discussion in *11*). The theory based on meiotic drive is discussed in the text. The possibility that the large X effect is caused by the misregulation of the X chromosome in the male germline (including the idea that Meiotic Sex Chromosome Inactivation is disrupted in hybrids) has not been theoretically worked out but repeatedly pointed out empirically (see discussion in *11*).

#### 3.2 Comparison of the regulatory theory with the dominance theory

The dominance theory for Haldane’s rule proposes that recessive incompatibilities occur, such that, in a hybrid F1 autosomal background, F1 females are fit while F1 males are unfit. The reason why F1 females are fit is that incompatibilities carried on the X are “recessive”, meaning that if the X were homozygous (in an F1 hybrid background), these incompatibilities would manifest themselves. If such females could be produced in a backcross or through genetic manipulations (such as the F1 females with attached X in *Drosophila*), they would therefore have low fitness. The theory further considers that hemizygous males express the incompatibilities because, having a single X, these incompatibilities carried on the X cannot be masked. The theory has to account for the fact that XX F1 females, having two X carry twice as many incompatibility genes compared to XY F1 males carrying a single X. Hence incompatibilities must be recessive on average for the theory to hold (*22*, *99*). Different models proposed different ways to achieve this pattern. For instance, different beneficial alleles can fix in different populations and some can turn out to be incompatible in the sense just explained.

In the model of stabilizing selection proposed by Barton (*24*), combinations of phenotypically dominant mutations can contribute to bring a population closer to an optimum. When different dominant mutations (on autosomes and on the X) coming from two parental species who adapted independently, combine in an F1 female, they all add-up to result in a near optimal phenotype (since all the mutations coming from each parental species independently add-up, as they are all dominant and have additive effects between loci). In contrast, F1 males miss the dominant mutations that occurred on the X of one parental species, causing a departure from the optimal phenotype (this model assumes no mutation on the degenerate Y). “Homozygous” F1 females would also miss all the mutations from the X of one of the two parental species (while having the dominant mutations on autosomes from both species), resulting in low fitness. Note that this model does not require specifically a different number of dominant mutations on the X of the two species.

Another model of stabilizing selection has been proposed that includes additive mutations (within and between loci), but assumes that mutations in males have twice the phenotypic effect when hemizygous (although not explicitly mentioning expression levels, this assumption might be viewed as a way to account for the occurrence of dosage compensation) (*25*, *26*). As in Barton’s model, F1 females remain close to the phenotypic optimum because all mutations are additive. F1 males, in contrast, depart from the optimal phenotype because the mutations on their X mismatch with half the additive autosomal alleles (coming from the same parental species as the X), while the other half of additive autosomal alleles do not add up with the missing X (from the other parental species). Here too, the “homozygous” females exhibit the same phenotype, and low fitness, as the F1 males.

All these models do not propose the same processes to explain HR. In particular, they consider different mutational input (the occurrence of recessive incompatibilities, the occurrence of phenotypically dominant mutations or the occurrence of additive mutations that have twice their phenotypic effect when hemizygous, respectively). They have one pattern in common, besides predicting HR: they all predict that a female that would be homozygous for a parental X in an otherwise F1 background has a lower fitness than a regular F1 female. This implies a dominance effect on the X, i.e., heterozygotes have a higher fitness than the average of the two homozygotes. In all cases the fitness of hemizygous F1 males is also found to be the same as the fitness of females made homozygous for the X. For this reason, and despite being based on different types of mutations, they can all be viewed as specific cases of a “general” dominance theory.

Our theory is based on another type of mutational input (additive mutations on cis and trans-regulators), but bears some similarity with these previous models in that HR is generated by a deficit in averaging between X and autosomal effects in the heterogametic sex relative to the homogametic sex. However, our theory does not fit well in the “general” framework of the dominance theory mentioned above. Indeed, in our regulatory model, F1 males show a reduced fitness compared to F1 females (Haldane’s rule), but the reason is not always a “recessive” effect, in the sense described above. Unlike in the dominance theory, the effect of the F1 genetic background differs between males and females: they express different trans-acting factors, both in the parental species and in the male and female hybrids (Fig S12). Hence, the fitness of hemizygous males is not necessarily the same as the fitness of “homozygous” females. Hemizygous males present the same fitness reduction as “homozygous” females only for genes located in ancestral strata (Eq. 14), but not in derived strata (§2.5, Fig S10c). This is particularly clear in the example shown in Figure S7a, where F1 females that would be made homozygous for the X would retain optimal expression, contrarily to F1 males. Generally, for derived strata, the disruption of dosage compensation always impacts more the heterogametic than the homogametic F1, although the effect varies with the type of dosage compensation and the position of the loci involved (on an ancestral or derived stratum, see Eq. 11). The difference between the fitness of F1 males and “homozygous” females also varies with type of dosage compensation and the position of loci, although “homozygous” females tend to have a lower fitness than F1 females (this is the case in the examples shown on Figures S7b and S8). Note too that in the case of derived strata, incompatibilities are, strictly speaking, not simple X-autosomes incompatibilities (see §2.5), as corresponding genes on the Y are not degenerate in one of the two parental species.

More fundamental differences between the “classical” dominance theory and our regulatory theory can be noted (table 1). First, our theory offers a mechanistic explanation of hybrid incompatibilities, specifically based on the misregulation of dosage compensated genes. Second, our theory does not simply postulate the existence of recessive incompatibilities (as if dominance was an intrinsic property of alleles). These incompatibilities emerge from the process of sex chromosome evolution and the coevolution of additive cis and trans-regulatory traits. These incompatibilities often turn out to be ‘recessive’, at least for dosage compensated genes in ancestral strata (see §2.3), but this is an evolved property mediated by the genetic background (the autosomal trans-regulators). A third difference is that our theory better explains why HR is based in sterility in some groups and viability in others (see §2.4). It also better explains why HR can sometimes be caused by W or Y-autosome interactions (see §2.5). Finally, it explains better HR on viability in species with X inactivation in females (see main text).

**Fig. S12.**
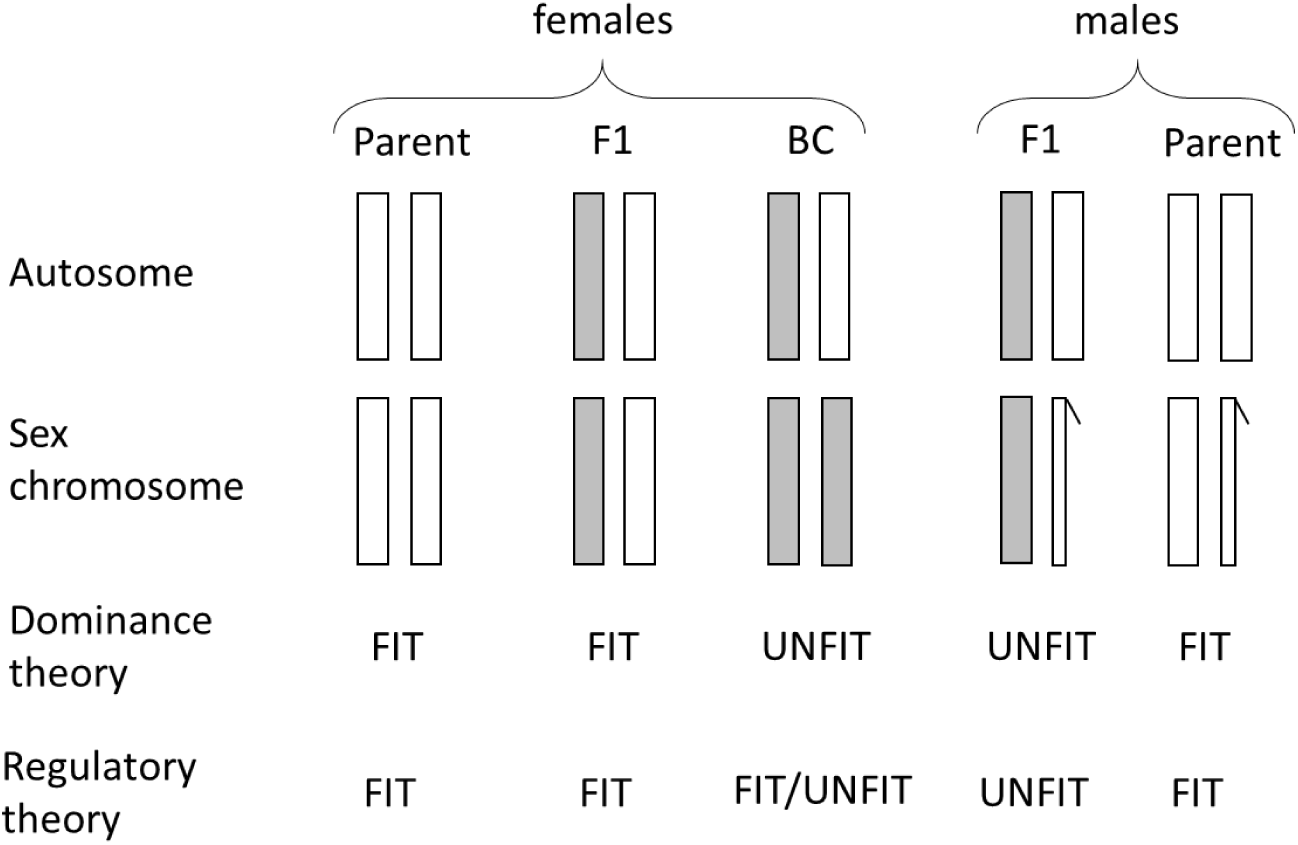
Simplified fitness patterns in the dominance versus regulatory theory. The chromosome with a hook represents the Y. A key difference is that the fitness of “homozygous” females differs between the two theories (indicated by “BC” as these females can be obtained by backcrossing or by creating “attached X” F1 females in *Drosophila*). In the dominance theory, “homozygous” females exhibit the same fitness reduction as F1 males. In the regulatory theory, male and female fitness can be uncoupled since they express different trans-regulators. The sex asymmetry results from lack of averaging of cis-regulator effects in males (see above).

#### 3.3 Criticisms of the dosage compensation hypothesis for Haldane’s rule

The list below presents past arguments against a theory of HR based on the disruption of dosage compensation. It has to be taken with precaution for three main reasons.

First, a firm theory of Haldane’s rule based on the disruption of DC has never been made, so most of these past arguments were made against a verbal and relatively vague theory. For instance, the relative level of expression disruption of autosomal vs. sex-linked genes in hybrids (as investigated e.g. in *86*, *95*) is sometimes mentioned as a decisive observation. Yet, this comparison is mostly uninformative. Autosomal misexpression reduces the fitness of both sexes in F1, and is therefore irrelevant to HR. It does not indicate whether the cause of HR is related to misexpression of sex-linked genes. If for instance DC is controlled by a global mechanism, such as in mammals or *Drosophila*, it might be constrained and slow evolving compared to autosomes. In this case, only few genes escaping this global DC might contribute to HR (including sex-limited genes in the germline not regulated by this somatic global DC). As a result, misregulation may appear larger overall on autosomes than on sex-linked genes, but this observation would not rule out that the cause of HR is misregulation of sex-linked genes.

Second, the idea that HR may be caused by DC disruption, made in the past, is much narrower than our regulatory theory. The idea usually considered is that the mechanism of dosage compensation is disrupted in hybrids, due to a divergence in the genes *directly* involved in the DC pathway. This is much narrower than in our regulatory theory, where the reduction of hybrid fitness is caused by the misregulation of gene expression *for all genes that are dosage compensated*, not only for genes directly involved in DC regulation. Our theory potentially includes most genes on the non-recombining part of the X or Z, not just the genes directly identified in the pathways controlling DC (the latter can nevertheless play a role in the misexpression of dosage compensated genes). If for instance, again, DC is controlled by a global mechanism, such as in mammals or *Drosophila* (although local DC probably precedes global DC, as seen on neo sex chromosomes (*130*)), only the genes escaping this global DC might contribute to HR, including sex-limited genes in the germline not regulated by somatic global DC. Their misexpression would play a key role in the fitness reduction of heterogametic F1, but they would be entirely unaffected by the global DC mechanism, and entirely unrelated to the control of this DC regulation. Note also that defining dosage compensation for sex-limited genes can only be made here relative to the ancestral expression level (before the evolution of recombination suppression). Since such genes are only expressed in one sex, male-female comparisons cannot be made, as usually done, to assess DC. In these cases, comparing expression levels in parental species and the hybrids would be important.

Third, the genetics of post-zygotic isolation is a broader topic than the genetic cause of HR. Many incompatibilities can impact hybrid fitness, in addition to those causing HR (i.e. not specifically reducing the fitness of the heterogametic sex). Importantly, such incompatibilites can be caused by many other mechanisms than expression disruption. Even in the context of HR, exceptions to the rule are known and relatively well understood (in particular nucleo-cytoplasmic incompatibilities (*75*)). Hence, assessing the merit of the different theories of HR based on a specific gene disrupting hybrid fitness in a specific cross must be done carefully. Documenting broad patterns such as those mentioned in Table 1 could be a more informative and more robust way of comparing the different theories.

With these caveats in mind, here are 13 arguments made in the literature against a (relatively narrow) theory based on the idea that the cause of HR is due to the direct disruption of the dosage compensation pathway in hybrids.

The 1^st^ major critique of the DC hypothesis is that the rule was observed in groups in which DC was allegedly absent (such as in birds and lepidopterans (*1*, *3*)). However, DC has since been documented in these groups (*59*, *61*), although global DC is perhaps less frequent in ZW species(*131*).

The 2^nd^ argument is based on an experiment with *Drosophila* (*52*). However, this experiment was based on crosses with mutants in which DC was supposedly fully functional, but without showing that this was the case. Hence, the results presented could not really discard DC disruption as a cause of HR. Specifically, it could not discard the possibility that hybrid male sterility was caused by a failure in DC downstream of *Sxl* regulation (which is the sex-determining switch in Drosophila).

The 3^rd^ argument is that DC evolves very slowly, being an essential function under strong constraints (*53*). However, as our results show, divergence in DC can readily occur, even with substantial stabilizing selection on expression levels. Note that DC evolution could be triggered by different phenomena, including coevolution with cytoplasmic bacteria targeting DC pathways to achieve male killing (*132–134*).

The 4^th^ argument proposes that if Haldane’s rule was caused by DC disruption, cis and trans regulators involved in DC should exhibit signs of divergence between species exhibiting Haldane’s rule. Yet, Jaffe and Laird (*135*) reported unpublished data where the X-linked *D. pseudoobscura Hsp82* gene remained dosage compensated when transformed into various autosomal sites in *D. melanogaster,* suggesting conservation of the cis-regulatory elements involved in DC for ∼20 million years. Indeed, if cis-regulators involved in DC are highly conserved, they could not cause Haldane’s rule (*1*, *52*). However, further evidence showed that this conclusion, besides applying only to *Drosophila*, was premature. More recent investigations revealed that both cis and trans regulators involved in DC were actually fast evolving in *Drosophila melanogaster*. This is the case with *msl*, *mof,* and *mle* trans-acting genes (*53*, *84*) as well as cis-acting binding sequences on the X (*54*). The case of male lethality in *D. melanogaster* x *D. simulans* hybrids is not clear-cut. Some studies support a role of DC in male lethality (*136*, *137*) while others challenge this interpretation (*138*). In this cross, both males and females show a reduction in viability, but to a different degree depending on temperature. However, the effect of mutants is not always evaluated in this GxE context. Barbash shows that deletion of mel-specific DC genes—ie, forcing Xmel/Ysim hybrid males to develop using only *simulans* DC complex components— does not exacerbate F1 male lethality when this lethality was first rescued by lhr mutation. The test is based on the premise that the lhr rescue occurs and that playing with DC genes should show an independent effect when manipulated. However, lhr might be acting by interacting with these DC genes, so there may be no clear ‘independent’ manipulation of the DC phenotype in the first place in the experiment. Barbash also argues that DC disruption is unlikely because a lower-than-expected number of genes downregulated in lethal F1 males are located on the X chromosome. However, only the misregulation associated with sex chromosomes will have a differential fitness effect on males vs females. HR depends on the presence of the latter, not on the proportion of misregulation of X versus autosomes. Whether there is more or less than “expected” misregulation on the X says nothing about the cause of HR. What can be noticed, however, although it could be a coincidence, is that major genes involved in these hybrid incompatibilities interact with the dosage compensation complex. For instance, *Lhr* interacts with HP1, a chromodomain-containing protein that localizes to heterochromatic regions of chromosomes (*139*) and is also involved in DC (*140*). Nuclear pore complex proteins also cause hybrid male lethality (*141*, *142*), and are also involved in DC (*143*).

The 5^th^ argument is a refinement of the fourth. After the accumulating evidence demonstrating fast DC evolution in *D. melanogaster*, Tang and Presgraves (*142*) suggested that this phenomenon was not general and limited to that species. They concluded that the limited evolution of the MSL complex in *D. simulans* could not well explain the rapid evolution of Nup160 and the other Nup107 subcomplex genes in that species (these autosomal simulans alleles being involved in male hybrid lethality through incompatibility with the melanogaster X). This may be the correct interpretation, and this specific case may be regarded as mere exception to the general case. However, this argument can be nuanced by several points, that might warrant further investigation. First, the *mof* gene involved in the DC complex does show evidence of rapid adaptive evolution in *simulans* (*84*). Second, Nup160 and the other Nup107 subcomplex genes are known to be involved in DC in *Drosophila* (*143*). Third, DC disruption is not necessarily limited to the coevolution of the MSL complex and its cis-binding sites. For instance, MSL cis-binding sites could change location on the X, thereby changing the pattern of DC, as shown by the rapid and extensive turnover of individual binding sites of roX lncRNAs, which are essential for *Drosophila* DC (*57*). Fourth, some genes are dosage-compensated but non-MSL-binding, suggesting that MSL is not the only mechanism for achieving DC (*104*). This is also particularly the case for genes expressed in the germline, impacting male fertility in *Drosophila* (*66*).

The 6^th^ argument, also by Tang and Presgraves (*142*) and also concerning Nup160 in *D. melanogaster*, note that Nup160sim kills both Xmel/Ysim hybrid males and Xmel ·Xmel/Ysim hybrid females, and in a similar way, suggesting a common cause. This would exclude the role of DC, on the premise that DC only concerns males in *Drosophila*. Like above, this may be the correct interpretation, and this case may well be an exception (there are several other known exceptions to HR in other groups, involving specific Y effects or incompatibilities with mitochondria, etc.). However, interestingly, our results show that, especially for ancient Y, unbalanced females should show an equal decrease in fitness to males, especially for viability (Eq. 14) (for hybrid sterility, it is easy to uncouple the fitness effect seen in males and unbalanced females: it is sufficient to have male-limited genes, as discussed above). This result relies on a quantitative DC trait divergence between species. In our model, the *Drosophila* DC system corresponds to a particular case where the strength of X cis-regulators does not change and where male trans regulators double X expression in males (case *t_m_* = 2 in our notation). Any quantitative departure from this point (where the strength of X cis regulator changes) involves a correction in females. In a quantitative model, DC is a phenomenon that always involves both sexes, as long as *t_m_* is not exactly 2. Hence, anytime males will show DC disruption (because of *t_m_* divergence), so will unbalanced females.

The 7^th^ argument has not directly been made against the DC hypothesis but against the dominance theory. However, this issue also concerns the DC disruption hypothesis. Haldane’s rule is observed in species lacking heteromorphic sex chromosomes. In particular, in the genus *Aedes*, Haldane’s rule applies to many interspecific crosses (*17*). Yet, *Aedes* have homomorphic sex chromosomes with a sex locus. A priori this observation would rule out theories based on hemizygosity (dominance theory) or DC. However, recent genomic data have revealed that in *Ae*. *aegyptii*, the sex “locus” is a 1.5 Mb region with 30 genes in a 100 Mb non-recombining region encompassing the centromere of chromosome 1, that diverged between males and females. Hence, the premise that these species have a sex chromosome with just a sex-determining locus seems erroneous. More work is required to evaluate the extent of hemizygosity and the occurrence of (local) DC in these species, but the conclusion favoring the “faster male” hypothesis based on these *Aedes* data certainly requires re-evaluation.

The 8^th^ argument is that one might expect a breakdown in dosage compensation to affect the homogametic more than the heterogametic sex in cases where DC involves an active mechanism in the homogametic sex, such as in mammals or *C. elegans* (*3*). This argument does not hold up theoretically. The larger impact of DC disruption in the heterogametic sex holds up irrespectively of the type of DC (Eq. 11). Despite showing extensive misregulation in *Caenorhabditis* hybrids, especially involving males, and involving X-autosome and cis-trans coevolution, a hallmark of DC disruption, Sánchez-Ramirez et al. (*47*) excluded DC as an explanation based on the observation that expression levels were not strongly disrupted in females. This reasoning is based on the idea that DC is a female phenomenon, as it works by halving X expression in females. This argument (similar to that of Laurie) does not hold, since it is in fact both a male and a female phenomenon (the halving in females corrects X overexpression evolving in both males and females). Even with a *C. elegans-*like system, DC disruption will be larger in males than in females (Eq. 11).

The 9^th^ argument is that DC could play a role, but only in the case of species with fast and ongoing Y degeneration, dramatically reducing the scope for the general application of the DC hypothesis (*51*). Indeed, the effect of DC disruption has been understood as resulting solely from the mismatch of having non-functional (but dosage-compensated) genes on the Y in one species but not in the other (*51*). However, this is an incomplete picture, and overly restrictive, as DC disruption may also occur for genes that are degenerate in both species (Eq. 7, Fig 11).

The 10^th^ argument is that the DC hypothesis is supposed to entail that “*any* anomaly (not just sterility and inviability) appearing in hybrids results from a disturbance in dosage compensation” (*1*). Since morphological anomalies in hybrids do not seem to be more severe in *Drosophila* hybrid males, it would then indicate that DC disturbance is not occurring (*1*). This argument seems greatly exaggerated, in the sense that there is no reason to suppose that all hybrid problems are DC-related. For many traits, the genes involved may not be sex-linked, so there is no reason to expect that they all show an HR pattern.

The 11^th^ argument is that DC disruption could hardly predict partial hybrid sterility or inviability (*1*). This would be true if DC was an all-or-nothing phenomenon, but, as our results show without ambiguity, a partial fitness reduction is very easy to obtain after short divergence. As is now better understood, DC occurs often on a gene-by-gene basis, even when a global DC system is in place (*59*, *144*). This observation was not available at the time this critique was formulated, and is thus, now, less convincing than it was.

The 12^th^ argument is that “it is difficult to envision how the failure of dosage compensation could explain the sterility of hybrids that are viable and morphologically normal” (*1*). In this view, disruption of dosage compensation should affect hybrid viability more than fertility (*14*). This argument is close to the previous one, in the sense that DC is viewed as a global chromosomal level process. However, it would be fairly easy to observe sterile but viable hybrids: it only requires that DC disruption only involves a few sterility genes in recently derived species. The same explanation applies (and was applied) to the dominance theory: it would only involve the occurrence of recessive incompatibilities on sterility genes in recently derived species (*99*). We also directly show how genes only expressed in the heterogametic sex (like genes involved in fertility) can produce a stronger fitness reduction in heterogametic F1, compared to genes expressed in both sexes.

The 13^th^ argument is that, in *Drosophila*, the absence of dosage compensation in the male germline excludes its disruption as a cause of hybrid male sterility (*63*). This argument does not take into account that some form of dosage compensation may be occurring in the germline, even if does not involve the global somatic DC mechanism. Some regulation seems to take place on both the X and Y in the germline, beyond the effect of MSCI (*66*, *73*). If this regulation is local (on a gene-by-gene basis) instead of following the global somatic mechanism, it would rather facilitate the evolution of regulatory divergence between species on those genes. We showed that male-limited genes could play a disproportionate effect on HR. This observation would tend to reinforce this conclusion. If DC divergence of viability genes is limited (because of the global somatic DC mechanism that remains constant), it would exacerbate the role of HR of genes expressed in the germline that escape this global regulation.

